# Catalytic mechanism of the mitochondrial methylenetetrahydrofolate dehydrogenase/cyclohydrolase (MTHFD2)

**DOI:** 10.1101/2021.11.03.467155

**Authors:** Li Na Zhao, Philipp Kaldis

**Affiliations:** Department of Clinical Sciences, Lund University, Box 50332, SE-202 13, Malmö, Sweden

**Keywords:** Folate-mediated one carbon pathway, MTHFD2 reaction mechanism, empirical valence bond (EVB), dehydrogenase reaction mechanism, cyclohydrolase reaction mechanism

## Abstract

Methylenetetrahydrofolate dehydrogenase/cyclohydrolase (MTHFD2) is a new drug target that is expressed in cancer cells but not in normal adult cells, which provides an Achilles heel to selectively kill cancer cells. Despite the availability of crystal structures of MTHFD2 in the inhibitor- and cofactor-bound forms, key information is missing due to technical limitations, including (a) the location of absolutely required Mg^2+^ ion, and (b) the substrate-bound form of MTHFD2. Using homology modeling and simulation studies, we propose that two magnesium ions are present at the active site whereby (i) Arg233, Asp225, and two water molecules coordinate Mg_A_, while Mg_A_ together with Arg233 stabilize the inorganic phosphate (P_*i*_); (ii) Asp168 and three water molecules coordinate Mg_B_, and Mg_B_ further stabilizes P_*i*_ by forming a hydrogen bond with two oxygens of P_*i*_; (iii) Arg201 directly coordinates the P_*i*_; and (iv) through three water-mediated interactions, Asp168 contributes to the positioning and stabilization of Mg_A_, Mg_B_ and P_*i*_. Our computational study at the empirical valence bond level allowed us to elucidate the detailed reaction mechanisms. We found that the dehydrogenase activity features a proton-coupled electron transfer with charge redistribution coupled to the reorganization of the surrounding water molecules which further facilitates the subsequent cyclohydrolase activity. The cyclohydrolase activity then drives the hydration of the imidazoline ring and the ring opening in a concerted way. Furthermore, we have uncovered that two key residues Ser197/Arg233 are key factors in determining the cofactor (NADP^+^/NAD^+^) preference of the dehydrogenase activity. Our work sheds new light on the structural and kinetic framework of MTHFD2, which will be helpful to design small molecule inhibitors that can be used for cancer therapy.

## INTRODUCTION

Chemotherapeutic drugs that aim to kill cancer cells unavoidably also kill healthy normal cells causing undesirable side effects. Methylenetetrahydrofolate dehydrogenase 2 (MTHFD2) has emerged as new drug target due to its expression only during embryonic development but not in adult tissue, and due to its amplification in cancer cells.^1–5^ Overexpression of MTHFD2 provides cancer cells with the necessary building blocks for nucleotide (purine and pyrimidine) biosynthesis during rapid proliferation.^2^ MTHFD2 is a novel anticancer therapeutic approach to target cancer cells without damaging healthy cells.^2,5-7^

MTHFD2, functions as homodimer, carries out both methylenetetrahydrofolate dehydrogenase (***D***) and cyclohydrolase (***C***) activities that are derived from its trifunctional precursor methylenetetrahydrofolate dehydrogenase, cyclohydrolase and formyltetrahydrofolate synthetase 1 (MTHFD1) through the loss of the *C*-terminal synthetase domain and a novel adaptation to NAD^+^ rather than NADP^+^ as cofactor for the dehydrogenase activity.^8^ Although the 5,10-methylenetetrahydrofolate (5,10-CH_2_-THF) dehydrogenase activity is well recognized as NAD^+^-dependent with an absolute requirement for Mg^2+^ and inorganic phosphate (P_*i*_) in NAD^+^ binding, the two ions have no effect on THF binding.^8^ Neither the cofactor (NADP^+^/NAD^+^) nor the ions are required for the MTHFD2 cyclohydrolase activity.^8^ Other enzymes with dehydrogenase activities such as its cytosolic counterparts MTHFD1, utilizes NADP^+^ as cofactor whereas the mitochondrial isozyme MTHFD2L, can use either NAD^+^ (when using NAD^+^ it also requires both Mg^2+^ and phosphate) or NADP^+^ for dehydrogenase activity.^7^ Recent studies suggest that MTHFD2 can achieve higher catalytic efficiency when using NAD^+^ rather than NADP^+^.^10^ When using 5,10-CH_2_-H_4_PteGlu_1_ and 5,10-CH_2_-H_4_PteGlu_5_ as substrates, the NAD^+^-dependent dehydrogenase activity was 8.5 and 2.4 times higher than its NADP^+^-dependent activity, respectively.^10^ The increase in maximal activity when using NAD^+^ as cofactor rather than NADP^+^ is important because it increases the production of formyl-THF in mitochondria,^11^ which meets the high demand for glycine and purine during proliferation in cancer and embryonic cells.^2^ However, which features of MTHFD2 contribute to the cofactor specificity change is currently unknown.

The monomeric MTHFD2 consists of D and C domains responsible for the ***D*** and ***C*** activities. The D/C domains of MTHFD2 form a cleft, which is composed of two *α/β* strands that assemble together. The walls of the cleft feature highly conserved residues.^12^ NAD^+^ and P_*i*_ are bound along one wall, while the substrate is bound at the interface between the two domains. MTHFD2 functions as a homodimer with homodimerization occuring by antiparallel interaction of the two NAD^+^-binding domains.^12,13^

Although magnesium has long been recognized as an essential metal cation for both the NAD^+^- and NADP^+^-dependent dehydrogenase activities of MTHFD2, it has been difficult to crystallize the MTHFD2 complex together with Mg^2+^.^11,14^ The monomeric crystal structure of MTHFD2 (5TC4) was solved at 1.89Å with NAD^+^, P_*i*_, and the inhibitor LY345899. Subsequently, homodimeric MTHFD2 was co-crystallized with three inhibitors (Compound 1, 6JID; Compound 18, 6KG2; and DS44960156, 6JIB) in the presence of cofactors (NAD^+^ and P_*i*_) at 2.25Å, respectively. However, the absolutely required Mg^2+^ is missing in all these 4 structures. Therefore, the location of Mg^2+^ in the MTHFD2 is not known.

P_*i*_ has been observed to occupy a position that is next to the 2’-hydroxyl of NAD^+^, which mimics the space that would otherwise be occupied by the 2’-phosphate of NADP^+^.^8^ Previous work has identified two residues, Arg201 and Arg233, that are important for the binding of P_*i*_ or the 2’-phosphate of NADP^+^.^8^ It is not known whether Mg^2+^ is directly involved in the P_*i*_ binding but the high Lewis acidity of Mg^2+^ is capable of stabilizing a phosphate anion. Characterization of MTHFD2 mutations at Asp168 and Asp225 indicate that both residues play a primary and direct role in assisting the binding and orientation of the Mg^2+^ ion in the cofactor binding site.^8^

Since MTHFD2 functions as dimer and no crystal structure of MTHFD2 with both inorganic phosphate and Mg^2+^ has yet been obtained, we have reconstructed the model of MTHFD2 homodimer complex based on available X-ray structures, site-directed mutagenesis and empirical valence bond (EVB) studies, and have quantum chemically located the Mg^2+^ binding sites.

To probe the energetics and mechanism of the initial proton-coupled electron transfer (PCET) process between NAD^+^ and 5,10-CH_2_-THF, and how it connects to the subsequent cyclohydrolase reaction, we have combined density functional theory (DFT) calculations with classical molecular dynamics (MD) simulations and hybrid quantum mechanics/molecular mechanics (QM/MM) calculations using the empirical valence bond approach. We propose a putative mechanism for the opening of the imidazoline ring after the PCET reaction and discuss its possible implications in driving the reaction into the desirable direction.

## MATERIALS and METHODS

### MTHFD2 Structure Construction

The initial structure for MTHFD2 was taken from Protein Data Bank (ID: 6KG2). 50 models were generated by “loopmodel” class within MODELLER 9.23^30^ to model and refine the missing loop from residues 281 to 285.^15^ [6R]-5,10-methylene-THF (endogenous one-carbon donor) was built from the ZINC database (ZINC4228244; https://zinc15.docking.org/substances/ZINC000004228244/; note that we have changed the D-glutamate moiety into its native L-glutamate in our simulation study; see Figure S2 and S3). Dock3.7 was used to dock 5,10-CH_2_-THF into all above 50 models. The conformations from the top ranked pose, where the reactive C atoms from 5,10-CH_2_-THF and NAD^+^ are within a catalytic feasible distance (4.5Å) and in catalytically competent orientation, were selected for further evaluation. In addition, we have added the absolutely required Mg^2+^ at the inorganic phosphate binding site after systematically examining the Mg^2+^ binding pockets in all known crystal structures and integrated this with the mutagenesis studies of MTHFD2 (the details are given in Supporting Material S1.2). The constructed MTHFD2 structure is shown in Figure 2. Key interactions between THF and MTHFD2 include (see Figure 2b): (i) the backbone of Val-131 and Leu-133, and the side chain of Asp-155, form hydrogen bonds with the Pterin group of THF; (ii) Tyr-84, *para-aminobenzoic* acid of THF, and the ring of nicotinamide of NAD^+^ form π-π stackings; (iii) Asn-87 forms a hydrogen bond with the glutamate moiety of THF. These interactions of THF with MTHFD2 recaptures the interaction scenario of inhibitor L34 in the 5TC4 crystal structure. The key interactions between NAD^+^ and MTHFD2 include (see Figure 2c): (i) the backbone of His-232 and Arg-233 form hydrogen bonds with the adenine group of NAD^+^; (ii) the backbone of Arg-201 and Ile-276, and side chain of Ser-202 and Asn-204 form hydrogen bonds with the ribose and phosphates moieties of NAD^+^; and (iii) the side chain of Thr-176, and the backbone of Val-274 form hydrogen bonds with the nicotinamide group of NAD^+^.

**Figure 1:**
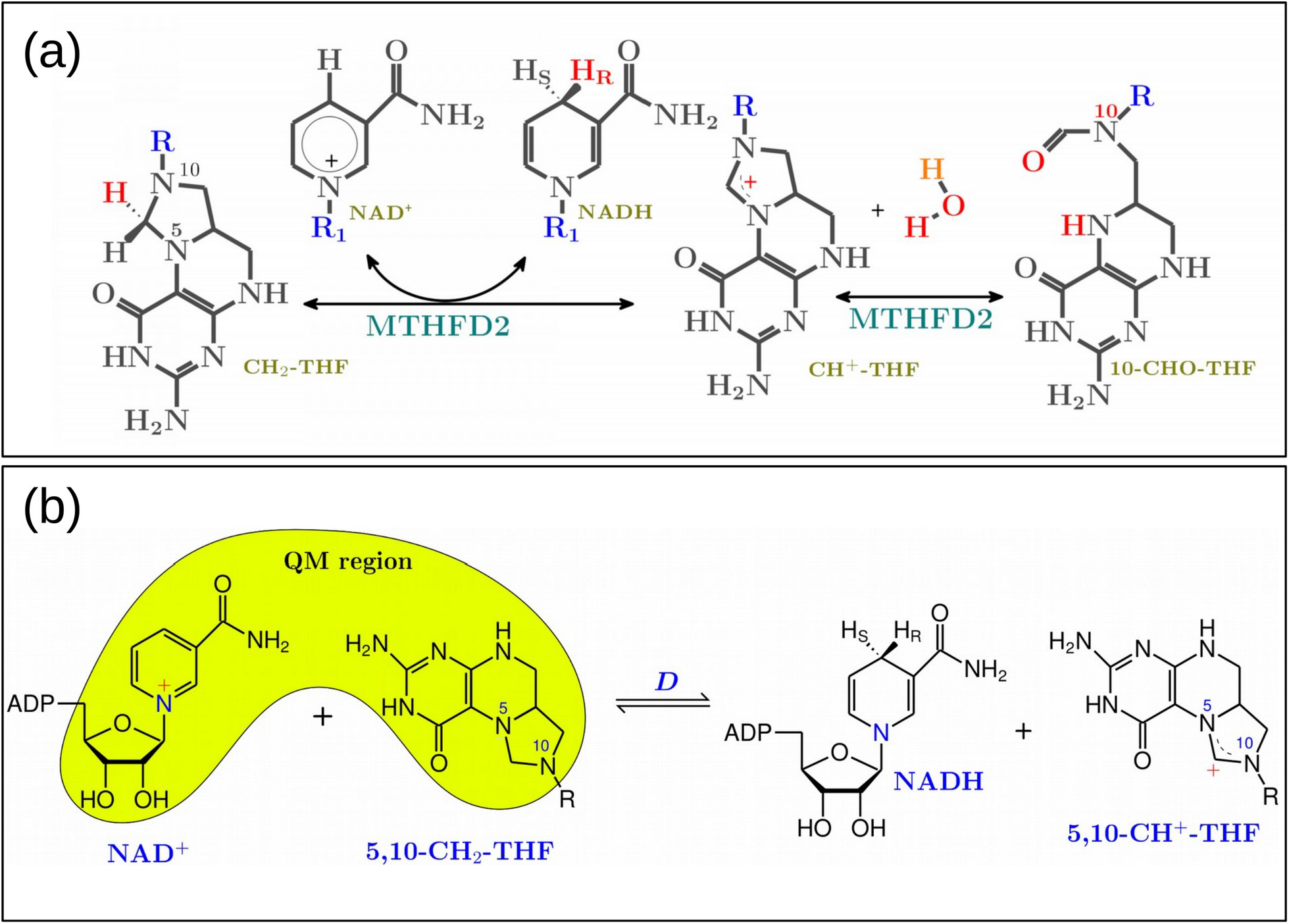
(a) The MTHFD2 reaction scheme. (b) The dehydrogenase (**D**) activity. Note that the QM region indicates the EVB atoms used for our empirical valence bond simulation of the **D** activity.

**Figure 2:**
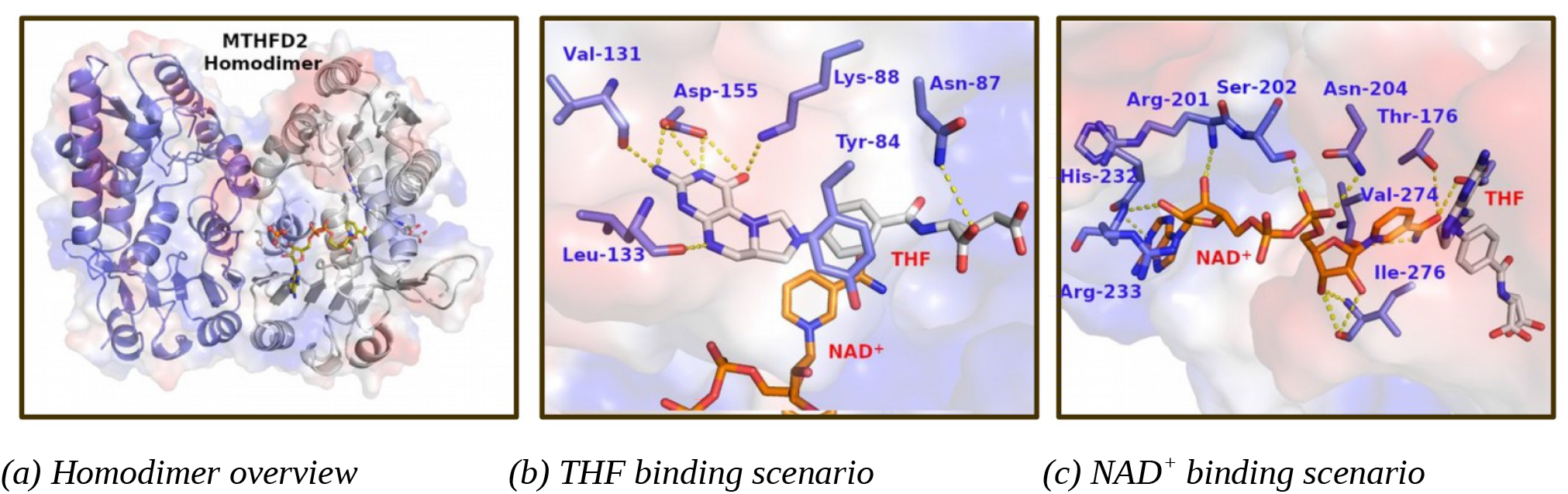
Overview of modeled system, with THF and NAD^+^ binding pockets.

### EVB Simulations

We have carried out density functional theory (DFT) calculations with a continuum solvent model to optimize the geometries of the reactant, transition state and product and the Mulliken charge for our EVB atoms using the B3LYP method with the 6-31g(d) basis set. In addition, we have used the B3LYP function with larger basis sets cc-pVTZ to obtain a kinetic description of the proton transfer from 5,10-CH_2_-THF to NAD^+^ with more accurate energies. The results from the DFT calculation (the active site model consisted of 38 atoms) were then used to calibrate empirical valence bond models for the proton transfer step. The two Mg^2+^ binding models were optimized in the MTHFD2·NAD^+^·P_*i*_·Mg^2+^·THF complex using constraint_pair within the Molarix-XG package to constrain by the force constant of 10.0 and around the distance that represents the configuration of the optimized Mg^2+^ configuration. The active site region along with the MTHFD2·NAD^+^·P_*i*_·Mg^2+^·THF complexes were immersed in a 32Å sphere of water molecules using the surface-constraint all-atom solvent (SCAAS) boundary condition.^16^ A 2Å layer of Langevin dipoles was applied outside of this 32Å region, followed by a bulk continuum. Atoms beyond the sphere were fixed at their initial positions and no electrostatic interactions from outside of the sphere were considered. The long-range electrostatics was treated with the local reaction field (LRF) method. The geometric center of the EVB reacting atoms was set as the center of the simulation sphere. The Monte Carlo proton transfer (MCPT) algorithm was used to optimize the charge distribution of all ionizable residues. MCPT simulates the proton transfer between charged residues in which the charge distribution is updated and evaluated with Monte Carlo approaches to identify the optimal charge distribution. Hence, the protonation state for the ionizable residues is determined by calculating MCPT. The EVB atoms are given in each section below. Detailed EVB simulation procedures are described in our previous work.^17–19^ All the EVB calculations were done using the Enzymix module within the MOLARIS-XG package.^16,20^

### The Water Flooding (WF) Approach

The determination of internal water molecules in enzymes, especially those at the active site with charges in protein interior, is a major challenge for simulation studies.^21^ The water flooding approach proved to be an efficient way to determine the most realistic configurations of water molecules by insertion of excess number of internal waters at first to over-saturate the protein, and then use the postprocessing Monte Carlo (MC) strategy to keep only the most likely configurations of the internal water molecules.^22^ Development, validation and application of the water flooding approach have been described extensively.^17,22,23^ Here we briefly list the key parameters we used in this study: In our simulation, the SCAAS surface constraints and the local reaction field (LRF) long-range treatment as well as polarizable ENZYMIX force field were used, and 10,000 steps of minimization followed by 200ps MD relaxation were done on our initial structure with a time step of 1.0 fs. The final structures were then used for WF simulations. During the WF simulations, a spherical hard wall was placed to prevent the exchange of the inside water molecules with the outside water molecules. The radius of the spherical hard wall for the cavity was set at 6.0 Å.

### Binding free energy calculations

The binding free energies of the substrate (CH_2_-THF) and cofactor (NAD^+^) at the presence of one magnesium ion (Mg_A_^2+^/Mg_B_^2+^) and two magnesium ions (both Mg_A_^2+^ and Mg_B_^2+^) were calculated by the semi-macroscopic version of the protein dipole Langevin dipole (PDLD) with the linear response approximation (PDLD-S/LRA). At first, we generated MTHFD2 complex configurations with the charged and uncharged forms of solute by carrying out explicit all-atom MD simulations with the surface-constrained all-atom solvent (SCAAS)^16^ and treating the long-range interaction with the local reaction field (LRF).^24^ Then we did the PDLD/S calculations on the generated configurations. The average value was used as the consistent estimation of the binding free energy. In total, we have generated 4 configurations for the charged and uncharged states, respectively. 2 ps run was done for each of these simulations at 300K. The dielectric constant is set 4 for neutral protein, and the effective dielectric constant is set 60 for the charge-charge interaction. The justification of our treatment concept has been discussed in great detail in Ref.25. This method has been established over decades to provide a reliable estimation of the binding free energies.^26^

## RESULTS

### The Tale of Two Mg^2+^ Cations

The only existing site-directed mutagenesis study for MTHFD2 was done in 2005 before publication of the human MTHFD2 homodimer structures.^8^ The residues that were targeted for mutagenesis experiments were Asp168 and Asp225. Asp216 is sandwiched between the two residues (see Figure 3a) but was not investigated.^8^ Both D168N and D225N affect Mg^2+^ binding directly^8^ and the minimum distance between Asp168 and Asp225 is 7.9Å, which is beyond the distance required to coordinate Mg^2+^. Furthermore, we observed from preliminary simulations for one Mg^2+^ ion that P_i_ is drifting away (data not shown), suggesting the need for a second Mg^2+^ ion. Therefore, we hypothesize that there are two Mg^2+^ present to coordinate the phosphate ion. Because of this, we modeled the system with two magnesium ions and relaxed it for 50ns with the detailed stable binding pocket for Mg_A_ and Mg_B_ shown in Figure 3b-c. Arg233, Asp225 and two water molecules coordinate Mg_A_, while Mg_A_ together with Arg233 stabilizes the two oxygens of P_*i*_. Asp168 and three water molecules coordinate Mg_B_, and Mg_B_ stabilizes P_*i*_ by forming hydrogen bonds with two oxygens of P_*i*_. Arg201 directly coordinates the P_*i*_. Furthermore, Asp216 contributes indirectly to positioning and stabilization of Mg_A_, Mg_B_ and P_*i*_, through three water-mediated interactions. Our data indicates that two Mg^2+^ ions are necessary for optimal organization of the active site of MTHFD2.

**Figure 3:**
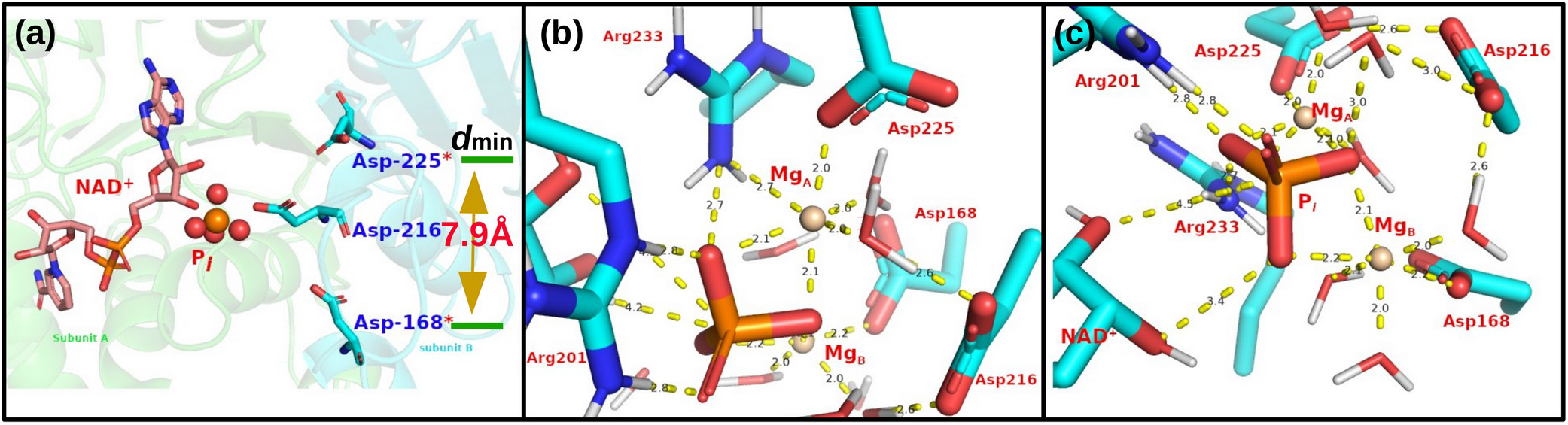
(a) NAD^+^ and P_*i*_ binding site from the X-ray structure (6KG2.pdb). Each monomer is labeled as subunit A (green) and B, respectively. Asp168, Asp216 and Asp225 from subunit B are colored as cyan for C atoms and shown as sticks. The minimum distance between Asp168 and Asp225 is 7.9Å. (b) The Mg_A_ binding pocket. (c) The Mg_B_ binding pocket.

### Enzymatic Catalysis

MTHFD2 combines (i) methylenetetrahydrofolate dehydrogenase and (ii) cyclohydrolase enzymatic activities. Early studies suggested that the Mg^2+^ and P_*i*_, ions bind to MTHFD2 first, followed by NAD^+^ and then 5,10-CH_2_-THF.^11,12^ The equilibrium ordered kinetic mechanism further indicated that 5,10-CH^+^-THF is released prior NADH, while in the NADP-dependent dehydrogenase reaction the same order of addition of substrate was suggested, but NADPH is released prior to 5,10-CH^+^-THF.^14^ The catalytic cycle of the NAD^+^-dependent dehydrogenase reaction can be summarized as six important intermediates: (1) the ternary complex (E:P_*i*_:2Mg^2+^); (2) the NAD^+^ quaternary complex (E:P_*i*_:2Mg^2+^:NAD^+^); (3) the Michaelis complex (E:P_*i*_:2Mg^2+^:NAD^+^:5,10-CH_2_-THF); (4) the quinary product complex (E:P_*i*_:2Mg^2+^:NADH:5,10-CH^+^-THF); (5) the NADH quaternary complex (E:P_*i*_:2Mg^2+^:NADH); and (6) the release of P_*i*_, Mg^2+^ and NADH. Note that both ***D*** and ***C*** activities share the same active site and the substrate can be channeled from ***D*** to ***C*** activity. It is unnecessary to release 5,10-CH^+^-THF, since it is required for the subsequent cyclohydrolase reaction. The integrated catalytic cycle including the cyclohydrolase activity is summarized in Figure 4.

**Figure 4:**
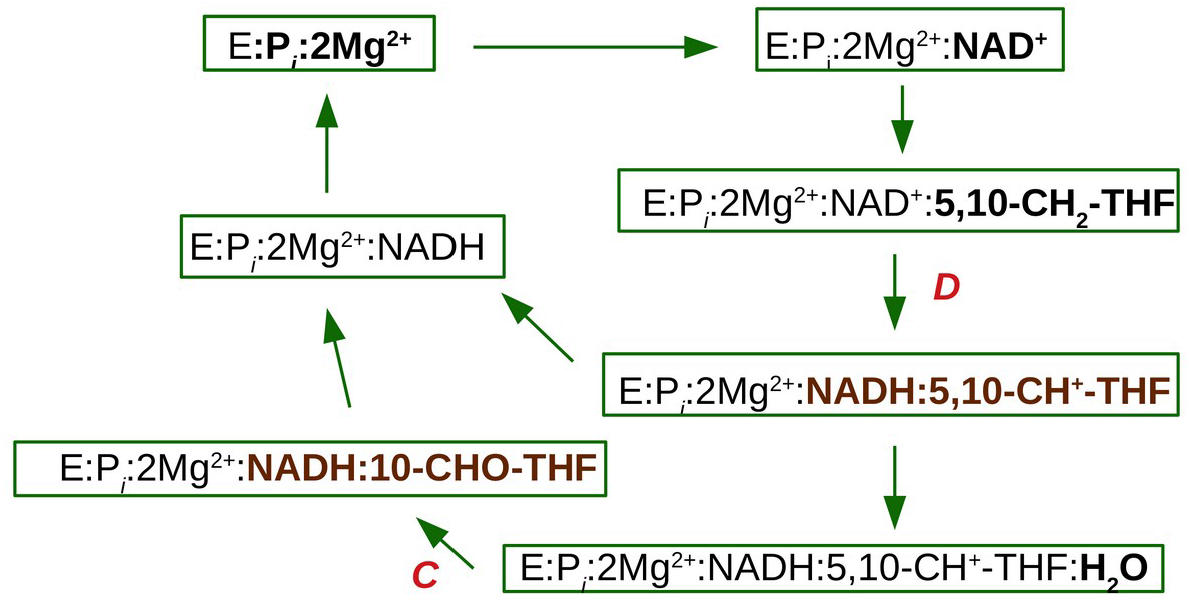
Catalytic cycle for the dehydrogenase (**D**) and cyclohydrolase (**C**) activities.

#### Dehydrogenase activity

The computational study of the dehydrogenase catalytic mechanism of MTHFD2 is described by three EVB resonance states (Φ_1_^D^, Φ_2_^D^, and Φ_3_^D^) that represent the reactant, intermediate, and product states (Figure 5). The hydride (H:) transfers from 5,10-CH_2_-THF to NAD^+^ in a *proton-coupled electron transfer* (PCET). This is electronically nonadiabatic with weak coupling between the reactant and product electronic states and ends with the electron delocalized in the pyridine ring of the nicotinamide moiety of the NADH. The reactive *C* atoms of THF and NAD^+^ in the optimized reactant geometry are at a 3.1Å distance from each other, with the transferring hydrogen at 2.1Å from the acceptor *C* atom. The π-π stacking interaction between the NAD^+^ pyridine ring of nicotinamide moiety and THF’s imidazoline ring connects the two reactants in the active site. The π-π stacking between Tyr84 and THF’s *p*ABA moiety orientating the THF binding and the flexibility of the (poly)glutamate tail of THF, was observed during simulations and can promote the formation of the transition state. In addition, PCET not merely involves substantial protein charge redistribution (from THF to NAD^+^) but also involves ordered solvent reorganization which is important for the subsequent reaction to proceed (Figure 5 and Figure S8).

**Figure 5:**
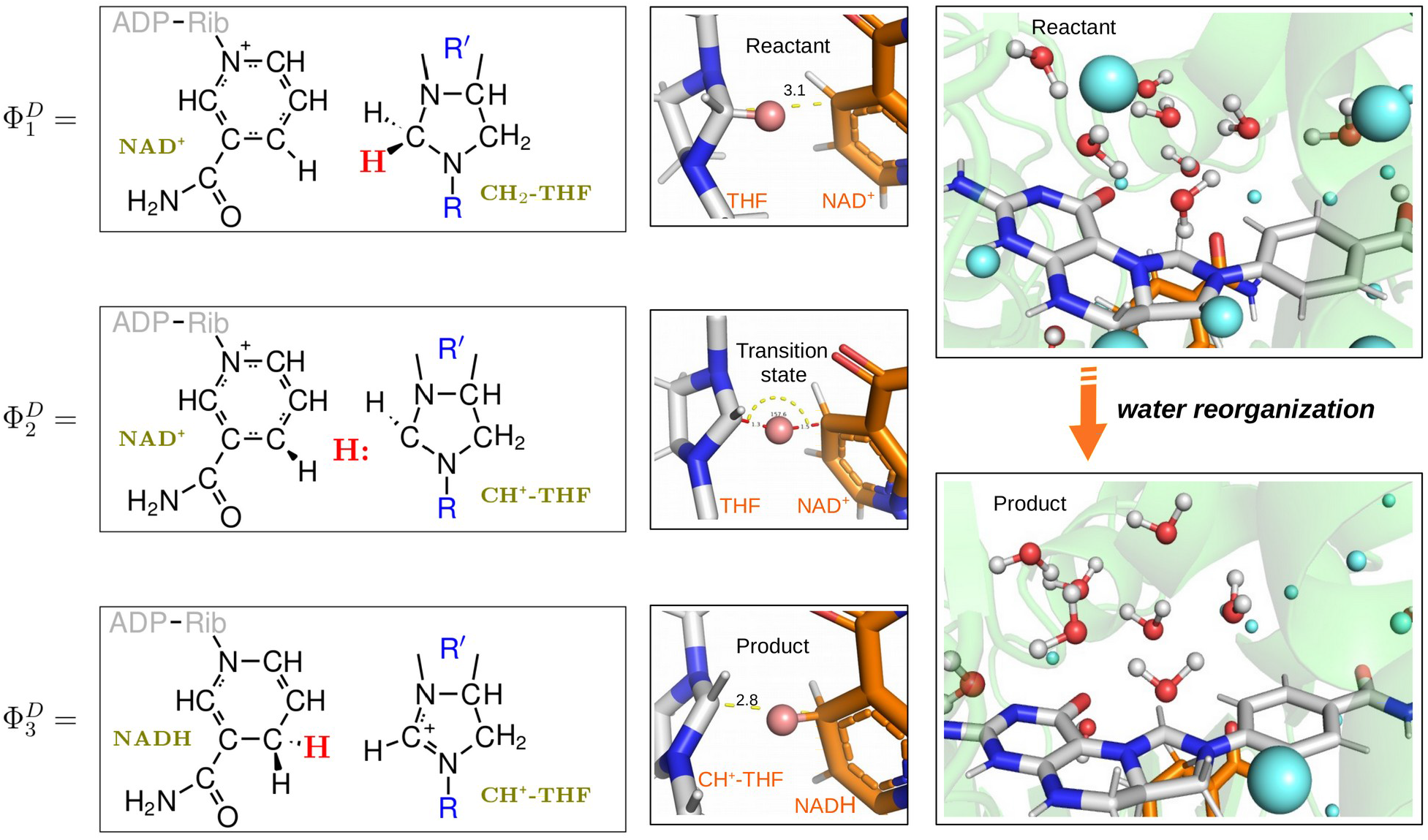
The left column shows the resonance states (Φ_1_^D^, Φ_2_^D^, and Φ_3_^D^) used to describe the dehydrogenase activity. The middle column shows the snapshots of the reactant, intermediate and product state, which are depicted as stick and colored by elements (C atoms: white for THF and orange for NAD^+^/NADH). The hydroxide is shown as a sphere and colored in salmon red. In the optimized reactant geometry, the distance between the reactive C atom of CH_2_-THF and the reactive C atom of NAD^+^ is 3.1Å, with the transferring hydrogen at 2.1Å from the acceptor C atom. The last column shows snapshots representing the water reorganization from reactant state to the product state. The water molecules within 7Å from the reactive C atom of THF are shown as both stick and sphere and colored red for oxygen atoms, while for the water molecules beyond 7Å are only shown as blue spheres to distinguish them.

We have calculated the binding energy of THF, NAD^+^, and PO_4_ in the presence of one magnesium (either Mg_A_ or Mg_B_), two magnesium, and no magnesium, respectively (Table 1). We found that the presence of magnesium does not affect THF binding but there was a moderate effect on NAD^+^ binding and a strong effect on PO_4_ binding. Our conclusion is that the presence of Mg^2+^ is important for the PO_4_ binding. We have observed an interesting result from our free energy calculations concerning the reaction kinetics. Since we have no experimental information of the reactions in solution, we have calibrated our EVB results with the reference water run using the B3LYP function with cc-pVTZ basis sets.^27^ The presence of one magnesium ion (Mg_B_) seems more energy favorable than the two magnesium system, which is an important insight because the two magnesium ions may contribute significantly to PO_4_ binding while slowing down the reaction kinetics.

**Table 1:** Calculated free energy (kcal/mol) for the dehydrogenase activity of MTHFD2. The ΔG^D,‡^_obs_ is calculated based on the experimental value k_cat_ = 12.4±0.71s^-1^ using transition-state theory.^10^

| Systems | $\Delta G^{D,\ddagger}$ | $\Delta G^D$ | $K_M^{\text{THF}}$ | $K_M^{\text{NAD}^+}$ | $K_M^{\text{PO}_4}$ |
| --- | --- | --- | --- | --- | --- |
| Water | 18.7 | 11.7 |  |  |  |
| MTHFD2 with 2 $\text{Mg}^{2+}$ | 24.1 | 4.7 | -0.50 | -10.45 | -97.56 |
| MTHFD2 with $\text{Mg}_A^{2+}$ | 22.5 | 13.4 | -0.74 | -0.22 | -37.66 |
| MTHFD2 with $\text{Mg}_B^{2+}$ | 16.9 | 1.9 | -0.34 | -5.72 | -42.67 |
| MTHFD2 without $\text{Mg}^{2+}$ | 26.5 | 0.3 | -0.82 | 5.37 | -8.53 |
| $\Delta G^{D,\ddagger}_{\text{obs}} = 16.0 \sim 16.1$ | | | | | |

#### Cyclohydrolase activity

Both MTHFD2 and MTHFD2L exhibit robust cyclohydrolase activity,^9,10^ which are independent of their dehydrogenase activity and NAD^+^ does not affect the reaction.^28^ Due to the relatively fast rate of the reaction and high absorbance of the substrate, it is hard to determine accurate *k_cat_* and *K_M_* values experimentally.^9^

The cyclohydrolase activity involves the nucleophilic addition of the hydroxide anion to the imidazoline moiety and the opening of the hydrated imidazoline ring. No mechanism has been proposed for this reaction, indicating a void of information that needs to be filled.

For the enzymatic catalysis, the opening of the imidazoline ring has three requirements: (i) activation of a water molecule for nucleophilic addition; (ii) the formation of a putative hydrated imidazoline intermediate; and (iii) a base to promote the scission of the central bond by abstraction of the hydrogen from the putative hydrated imidazoline intermediate. Even though the nature of the protein active site cavity is an aprotic environment, the Lys88 side chain provides polar interactions that anchors the oxygen of the pterin group from 5,10-CH^+^-THF by direct hydrogen bonding, and further interacts with the side chain of Gln132. However, neither Lys88 nor Gln132 can be the base to promote the scission of the central bond by withdrawing the hydrogen from the intermediate (Figure S7). Here, we used the water flooding approach to estimate the number of water molecules that can be present at the active site. The number of water molecules, with an average of 8 water molecules, located right at the active site is significant enough to form a protic proton-recycle system (see Figure 5 and Figure S8) that could facilitate a low-barrier hydrogen transfer/exchange. The possibility of forming a proton-relay system that recycles/exchanges the hydrogen to favor the cyclohydrolase reactions is tantalizing because it resolves which base abstracts the hydrogen from the hydrated imidazoline intermediate. Using the hydrated active site and computational approaches, we observed a concerted reaction mechanism for the hydration of the imidazoline ring (Figure 6). One hydrogen from water was abstracted by the pyridine-nitrogen atom of the imidazoline ring, while the hydroxide of the water molecule was attracted to the pyrrole-nitrogen atom. Since the pH inside the mitochondria is slightly above 7 (range from 7.0 to 7.8),^29^ this alkaline environment may facilitate the abstraction of the hydrogen from hydroxide to facilitate the scission of the central bond.

**Figure 6:**
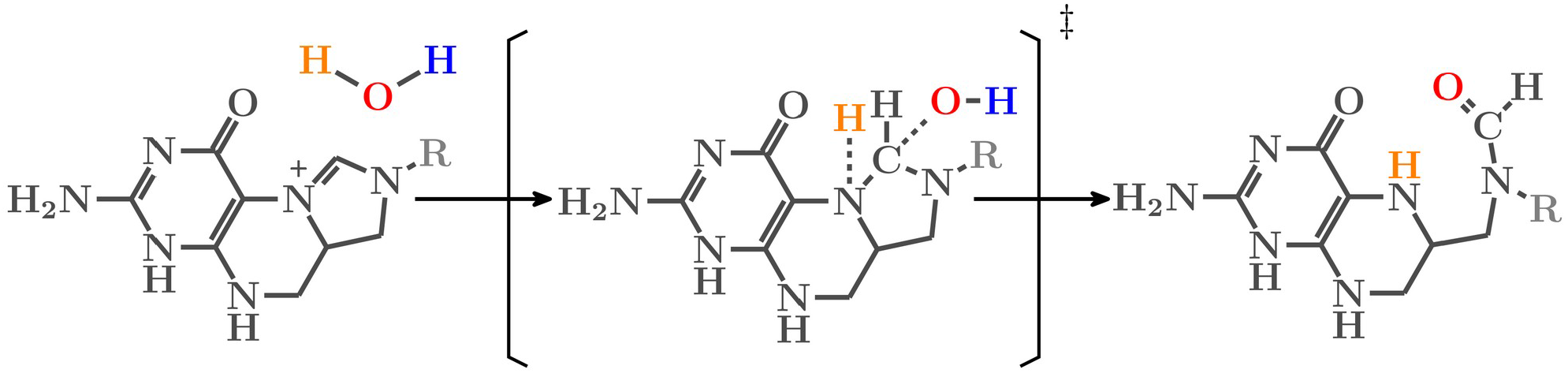
Proposed cyclohydrolase reaction mechanism.

**Figure 7:**
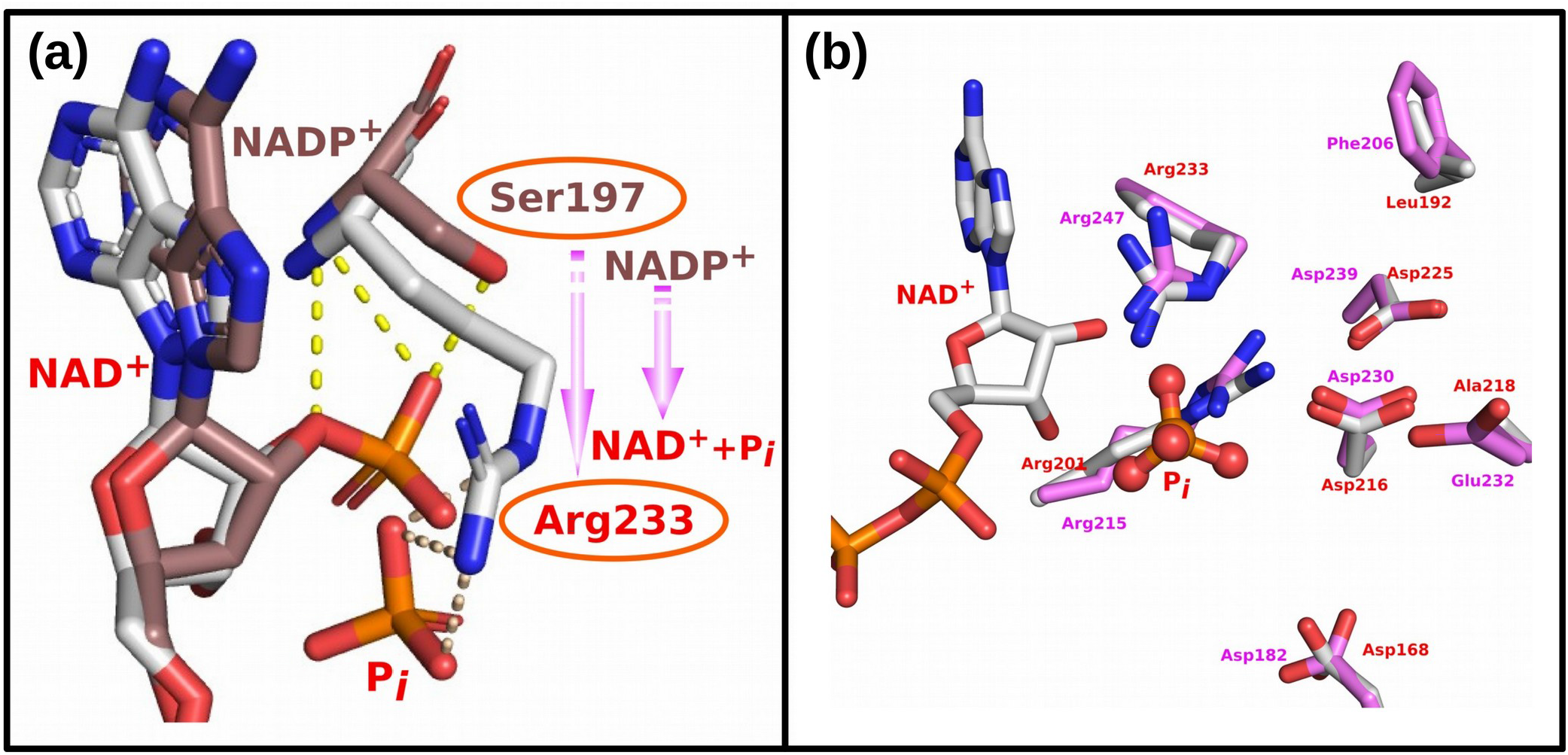
(a) Superimposition of the NAD(P)^+^ binding pockets with regards to their dehydrogenase activity. The C atoms of MTHFD1 (PDB: 6ECQ) were colored in brown, and those of MTHFD2 (PDB: 6KG2) were colored in white. Switching from Ser197 to Arg233 defines a key force in binding 2’-phosphate of NADP^+^ rather than phosphate ion (P_*i*_). (b) Superimposition of the MTHFD2 and MTHFD2L NAD^+^ binding pockets. Key residues involved in binding are shown as sticks and colored red for MTHFD2 and pink for MTHFD2L.

Furthermore, it would not be catalytically “wise” to release the 5,10-CH^+^-THF immediately after the dehydrogenase reaction is completed since it needs to be at the same active site for the subsequent cyclohydrolase activity given that a single site is used for both dehydrogenase and cyclohydrolase activities.^11^ Hence, we suggest that 5,10-CH^+^-THF is channeled from the previous dehydrogenase reaction to the subsequent cyclohydrolase reaction, which is also supported by a study of its cytosolic counterpart MTHFD1.^30^ A conformational change, particularly the solute reorganization during the dehydrogenase activity, may be responsible for the activation of the cyclohydrolase process. Such a mechanism would increase the efficiency of MTHFD2.

### To be or not to be NAD^+^-dependent

The cofactor preference of MTHFD2 for NAD^+^ (rather than NADP^+^) in the dehydrogenase activity is believed to promote “a more thermodynamically favorable pathway to balance the pools of 10-formyl-THF during development”.^31^ Sequence and structure comparison of MTHFD1 and MTHFD2 (Figure S5) indicated that Ser197 is a key factor in binding the 2’-phosphate of NADP^+^, while Arg233 is important to bind the phosphate ion. The change from Ser197 in MTHFD1 to Arg233 in MTHFD2, enables switching the dehydrogenase activity from NADP^+^-dependent to NAD^+^-dependent. Mutagenesis of MTHFD2 indicates that the Arg233Ser mutant almost completely lost the NAD^+^-dependent dehydrogenase specific activity with a 50% decrease in NADP^+^-dependent dehydrogenase specific activity.^32^ Furthermore, since MTHFD2L has been reported to use either NAD^+^ or NADP^+^ as cofactor, we have used homology modeling to build the MTHFD2L structure and superimposed it with MTHFD2 (for sequence alignment see Figure S5). Residues near the NAD^+^ binding pocket are well-conserved, while more distant residues vary, which contributes to the difference in catalytic efficiency between the MTHFD2 and MTHFD2L. Since earlier kinetic studies have shown that the ions bind MTHFD2 before NAD^+^,^11,14^ the binding of ions also selectively prevents NADP^+^ binding due to the steric occupation of the 2’-phosphate group of NADP^+^ by ions.

### Experimentally evidence of the two Mg^2+^ system and its implications for catalysis

Nicotinamide adenine dinucleotide (NAD^+^), as an essential and ubiquitous coenzyme in metabolism, participates in fundamental and vital pathways including energy metabolism regulation, DNA damage repair and recombination, and post-translational modifications.^33,34^

In order to identify crystallized structures of NAD^+^ bound with Mg^2+^, we have extracted 31,477 structures from Protein Data Bank by searching for ‘NAD’ and ‘Mg’ as ligands. Among these, 142 structures contain both NAD^+^ and Mg^2+^ which are listed in Table S4. After scrutinizing these structures, we decided to focus on the crystal structure of two proteins: (1) ketol-acid reductoisomerase (PDB ID: 4KQX), and (2) domain 1 of NAD^+^ riboswitch with nicotinamide adenine dinucleotide (NAD^+^) and U1A protein (PDB ID: 7D7V, 7D7W and 7D81), the latter has two Mg^2+^ in close proximity of NAD^+^. However, the NAD^+^ does not interact with protein in the riboswitch structures, namely 7D7V, 7D7W and 7D81 (see Figure S5). Therefore, we will further investigate the structure of the ketol-acid reductoisomerase (PDB ID: 4KQX). The insights we gained from the structure of 4KQX (Figure 8), indicated the existence of two Mg^2+^ ions, which further strengthen our concept of two Mg^2+^ binding modes corroborating previous data from direct mutagenesis and QM/MM studies. However, our free energy calculations indicated that one magnesium ion shows a slightly more energy favorable reaction kinetics but in the terms of PO4 binding, the two magnesium system is more favorable.

**Figure 8:**
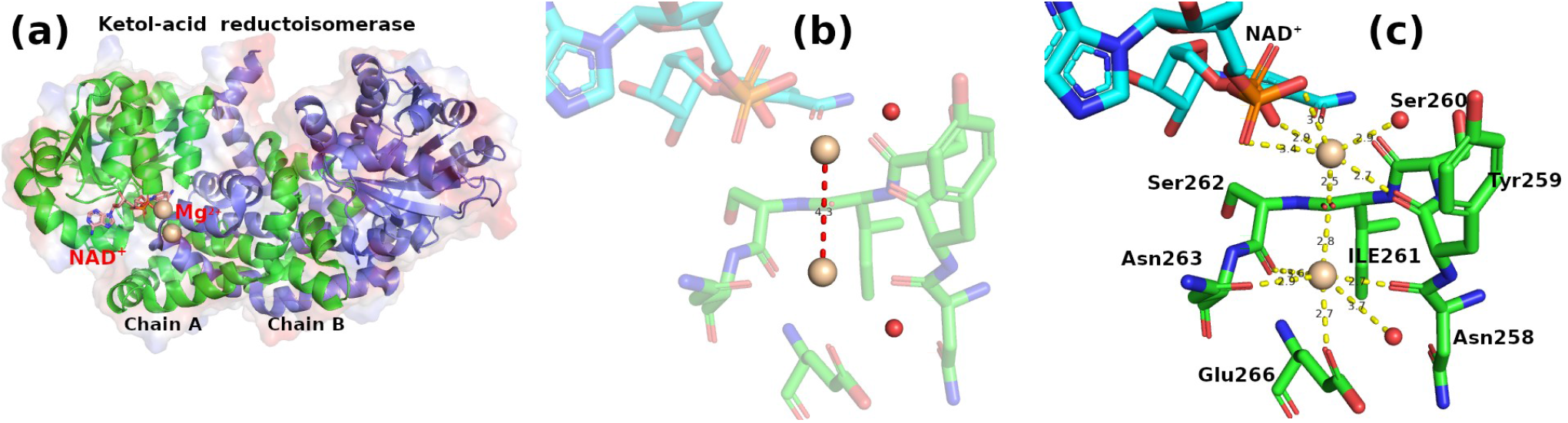
(a) Overview of the structure of Ketol-acid reductoisomerase (PDB ID: 4KQX). (b) The distance between two Mg^2+^ ions is 4.3Å. (c) The residues that coordinate the two Mg^2+^ ions system.

Magnesium exists as Mg^2+^ ion in the living system and plays an important, sometimes essential role in metabolic pathways and nucleic acid biochemistry. Notably the DNA polymerase features two key magnesium ions that facilitate the addition of new nucleotides to a growing DNA chain.^35^ Because we propose a two magnesium system from our modeling, our evaluation of the two magnesium system compared with a one/no magnesium system indicates that the presence of two magnesium significantly contribute to the PO_4_ binding but slightly slow down the reaction kinetics. Consistent with this, a study reported the pros and cons using two Mg^2+^ ions in the CDK2 reaction, where it can accelerate the reaction while slowing down the product release.^36^

### Implication for the drug discovery

As an ideal target for anticancer drug development, we need a drug highly specifically targeting MTHFD2,^5^ without affecting its cytosolic homolog MTHFD1 and the mitochondrial homolog MTHFD2L. Because MTHFD1 is expressed in normal tissues and most abundantly in the liver,^37^ inhibition of MTHFD1 would cause most likely unwanted side effects. MTHFD2L, sharing 60-65% identity with MTHFD2 and with a highly conserved substrate and cofactor binding pocket, is expressed in all tissues examined, with the highest expression levels in brain and lung in both humans and rodents.^38,39^ Therefore, a selective inhibitor for MTHFD2 over MTHFD1 and MTHFD2L is desirable as a lead for drug discovery to warrant successful clinical investigations. There are two apparent pockets in MTHFD2 for structure-based drug discovery: one is the cofactor (NAD^+^) pocket and the other is the substrate (CH_2_-THF) pocket. Since we know that NAD(P)^+^-dependent enzymes are ubiquitously expressed in metabolism and other cellular processes,^40^ inhibition of NAD(P)^+^ may induce significant toxicity. In addition, the substrate pocket is also present in other enzymes of the THF-mediated one-carbon metabolic pathway such as SHMT1/2, MTHFD1 and MTHFD2L. Therefore, interventions in the THF-mediated pathway in normal cells could induce undesired side effects. Hence, our study has highlighted a novel integrated pocket, different from the current cofactor and substrate clefts, which is formed at the interface between two monomers during dimerization (illustrated by balls in Figure S4). By exploiting this structurally less conserved allosteric site of MTHFD2, we could develop new inhibitors to help cancer patients. And a latest effort has been made into this direction with the xanthine derivatives being report at the allosteric binding site of MTHFD2, which prevents the binding of the cofactor and phosphate.^41^ Our preliminary screening has identified 3 clusters of compounds (see Table S1) that have potential to bind this structurally less conserved allosteric site. They neither fully occupy the substrates nor the cofactor binding site, but may protrude to the above two canonical binding sites which prevents the binding of either substrate or cofactor; it may coexist with the substrate/cofactor. Further structural and in vivo studies are required to validate these proposed compounds, thus to elucidate the mechanism of inhibition and further develop into candidate drugs.

## DISCUSSION

Even though it has been well recognized that there is an absolute requirement for Mg^2+^ in the MTHFD2 dehydrogenase activity, it has been a challenge to determine the location of the Mg^2+^ by X-ray due to the extremely low electron density and the small size.^42^ Scrutiny of the crystal structure of MTHFD2 indicates that the closest distance between Asp168 and Asp225 is 7.9Å (Figure 3a), and another residue Asp216 is sandwiched between Asp168 and Asp225 (PDB ID: 6KG2), which leads us to propose a model of two Mg^2+^ assisting in P_*i*_ binding. This model is further supported by the fact that one Mg^2+^ alone is not capable of stabilizing the P_*i*_ in our detailed QM/MM study. Our work has fully captured the presence of two Mg^2+^ cations and provided a clear picture of how they coordinate and stabilize the inorganic phosphate further contributing to catalysis. Furthermore, Asp168, Asp216, and Asp225 from chain A, and Arg201 and Arg233 from chain B, collectively coordinate the two Mg^2+^ and P_*i*_, which explains why MTHFD2 functions as a homodimer and homodimerization occurs by antiparallel interaction of the two NAD^+^-binding domains.

The dehydrogenase activity transfers two electrons coupled to the proton (H:) and this involves both electron relocalization featuring substantial protein charge redistribution and ordered solvent reorganization which activates the subsequent cyclohydrolase reaction. The cyclohydrolase activity features the opening of the nucleophilic attacked imidazoline ring, which is facilitated by the proton-recycling-network present at an aprotic pocket enabling low-barrier proton exchange. We were able to show that the Ser233 to Arg switch determines the cofactor preference from NAD^+^ to NADP^+^.

MTHFD2 is an important drug target lacking inhibitors in clinical trials or with FDA-approval. Our detailed structural analysis and mechanisms studies have uncovered (a) the presence and location of two Mg^2+^ ions; (b) the dehydrogenase activity orchestrated a proton-coupled electron transfer not only facilitating the proton transfer but also promoting the subsequent cyclohydrolase reaction; (c) a putative cyclohydrolase reaction mechanism; and (d) the key factors that determine the cofactor preference. Our study supports the importance of Mg^2+^ in the enzymatic activity of MTHFD2. In order to better understand the roles of two Mg^2+^ ions system, we dissected it into two interconnected roles: one is the structural role featuring the binding energy of substrate and co-factors and found it significantly contributes to the stabilization of the cofactor PO_4_; the other one is the functional role, where our study indicates that the disadvantage of two magnesium ions because it slightly slows down the reaction kinetics. Our detailed structural information has located the position of the two magnesium ions in MTHFD2, which provides a framework for the future characterization of the roles, the order and mode of binding of these two Mg^2+^ ions, as well as the underlying mechanism. In addition, the reaction mechanism has been investigated in an integrated way, rather than as a combination of isolated reaction steps, which may shed new light on this drug target.

Our work not only contributes to the understanding of the binding mode and catalytic mechanism of a drug target (MTHFD2), which is relevant not only for further studies on drug discovery, but also new cues concerning the coupling of metal and cofactor binding in the catalytic steps of other enzymes.

## ACKNOWLEDGMENTS

This work is supported by IngaBritt och Arne Lundbergs Forskningsstiftelse LU2020-0013 and the Crafoord Foundation (Ref. No. 20210516) provided for L.N.Z. P.K is supported by the Swedish Research Council (VR 2021-01331); Faculty of Medicine, Lund University; the Swedish Foundation for Strategic Research Dnr IRC15-0067; and Strategic Research Area EXODIAB (Dnr 2009–1039). Additionally, L.N.Z acknowledges the computational support provided by Prof. Warshel’s laboratory at the University of Southern California. Special thanks go to Prof. Ulf Ryde from Lund University who advised and commented on our manuscript, and Dr. Zhen T. Chu and Dr. Vesselin Kolev from USC. L.N.Z would love to thank Prof. Patrik Midlöv for his support in her research.

## Compliance with ethical standards

Ethical approval is not required for the study.

## Abbreviations

CH_2_-THF: 5,10-Methylene-tetrahydrofolate
CH^+^-THF: 5,10-Methenyl-tetrahydrofolate
CHO-THF: 10-Formyl-tetrahydrofolate
CH_3_-THF: 5-methyltetrahydrofolate
*C*: CH+-THF cyclohydrolase
*D*: CH_2_-THF dehydrogenase
EVB: empirical valence bond
LRF: local reaction field
MCPT: Monte Carlo proton transfer
MTHFD1: Methylenetetrahydrofolate dehydrogenase 1
MTHFD2: Methylenetetrahydrofolate dehydrogenase
MTHFD2L: Methylenetetrahydrofolate dehydrogenase (NADP^+^-dependent) 2 like
NAD^+^: Nicotinamide adenine dinucleotide
NADH: Nicotinamide adenine dinucleotide (NAD)^+^ hydrogen (H)
NADP^+^: Nicotinamide adenine dinucleotide phosphate
PDLD: protein dipole Langevin dipole
PCET: proton-coupled electron transfer
*S*: CHO-THF synthetase
THF: Tetrahydrofolate

## Modeled systems

### Structure background

In 2016, the first crystallization of MTHFD2 in complex with NAD^+^, inorganic phosphate (Pi), and LY345899 was solved at 1.89Å and provides a reliable template (5TC4.pdb) to start structure based inhibitor studies.^1^ The latest structure of MTHFD2 in complex with DS44960156 (6JIB.pdb), Compound 1 (6JID.pdb), and DS18561882 (6KG2.pdb) as well as cofactors (NAD^+^ and P_*i*_) were solved at 2.25Å and bears similarities with the first identified structure. With all these structures, the exact location of Mg^2+^ and its molecular interactions are still not known. Since MTHFD2 functions as dimer, we have reconstituted the active site of the MTHFD2 in a homodimeric complex (MTHFD2·NAD·P_*i*_·Mg^2+^·P_*i*_·THF) with inorganic phosphate, NAD^+^, and substrate using homology modeling.

**Figure S1:**
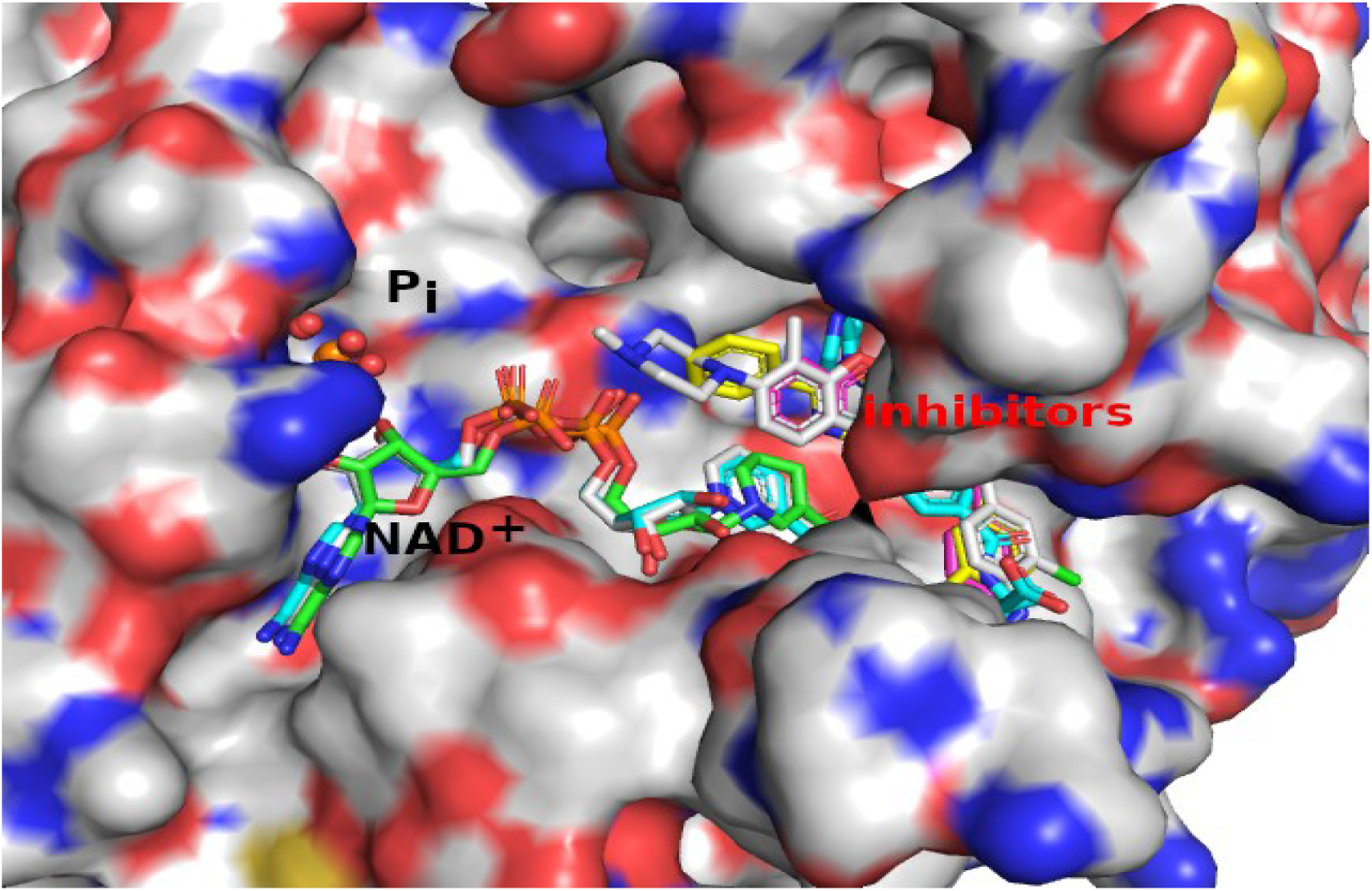
The superimposed snapshots of all crystallized MTHFD2 structures (1ZN4, 5TC4, 6JIB, 6JID, and 6KG2). The proteins are shown as surface. NAD^+^ and inhibitors are shown as sticks. Phosphate is shown as spheres.

### Construction of the two magnesium system

In order to investigate the crystallized structure of NAD^+^, PO_4_ bound with Mg^2+^, we have searched Protein Data Bank and 5J33.pdb was used for our initial modeling of the first Mg^2+^ in the MTHFD2 binding pocket, in which MTHFD2 is modeled using 6KG2.pdb as template. Based on the knowledge we have gained from mutagenesis (D168A/E/N/S, S201R, D225A/N and R233A/K/S), we have manually added the second magnesium and then optimized on several levels: first we optimized the binding pocket using ab initio and then we integrated the optimized binding pocket into the MTHFD2 complex. Several rounds of relaxation were carried out to let the system reach the equilibrium.

**Figure S2:**
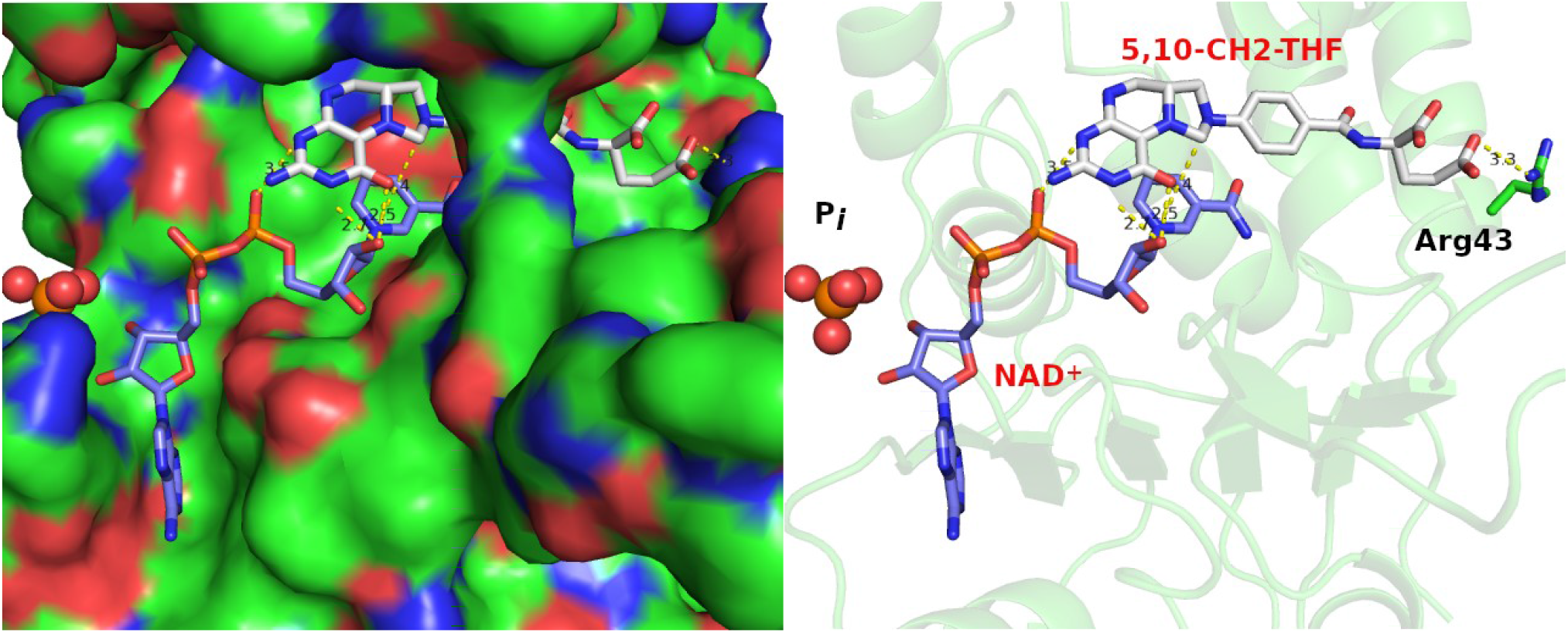
Representative docked pose with 5,10-CH_2_-THF and NAD^+^ in a catalytic feasible conformation. MTA, 5,10-CH2-THF and Arg43 are shown as sticks. P_i_ is shown as spheres.

**Figure S3:**
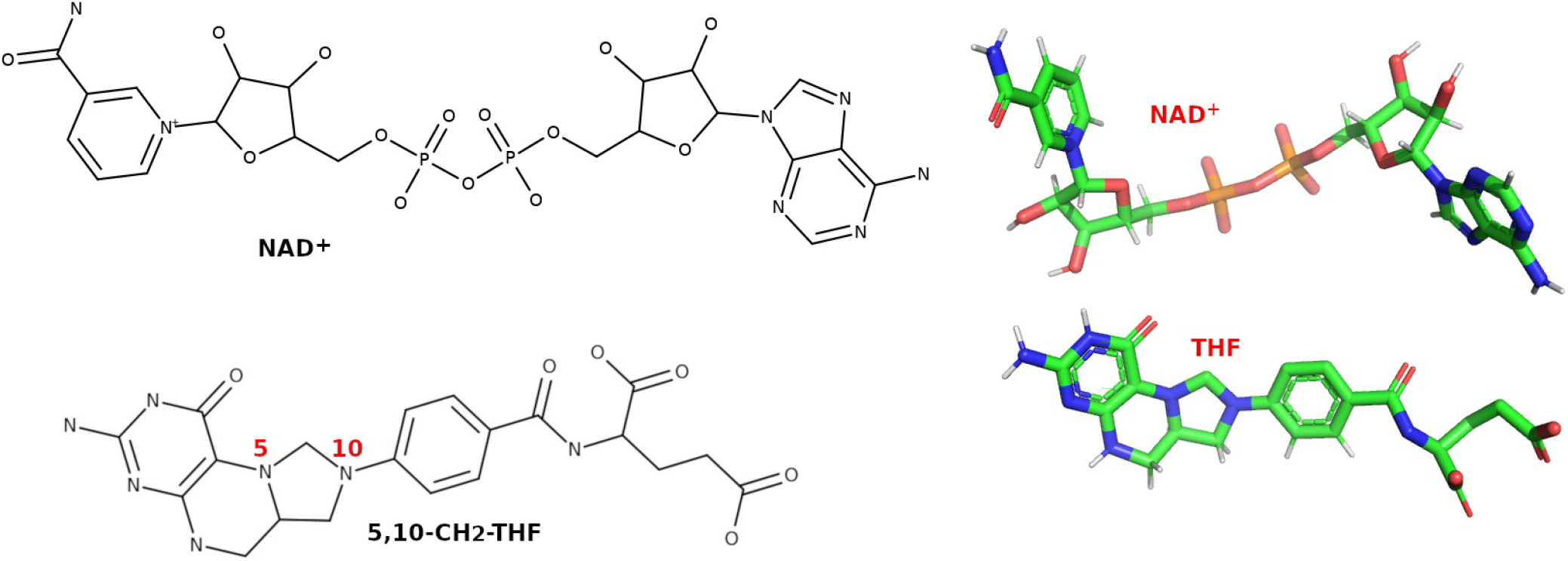
The chemical structure of NAD^+^ and 5,10-CH_2_-THF.

### Proposed homodimer binding groove and potential inhibitors

**Figure S4:**
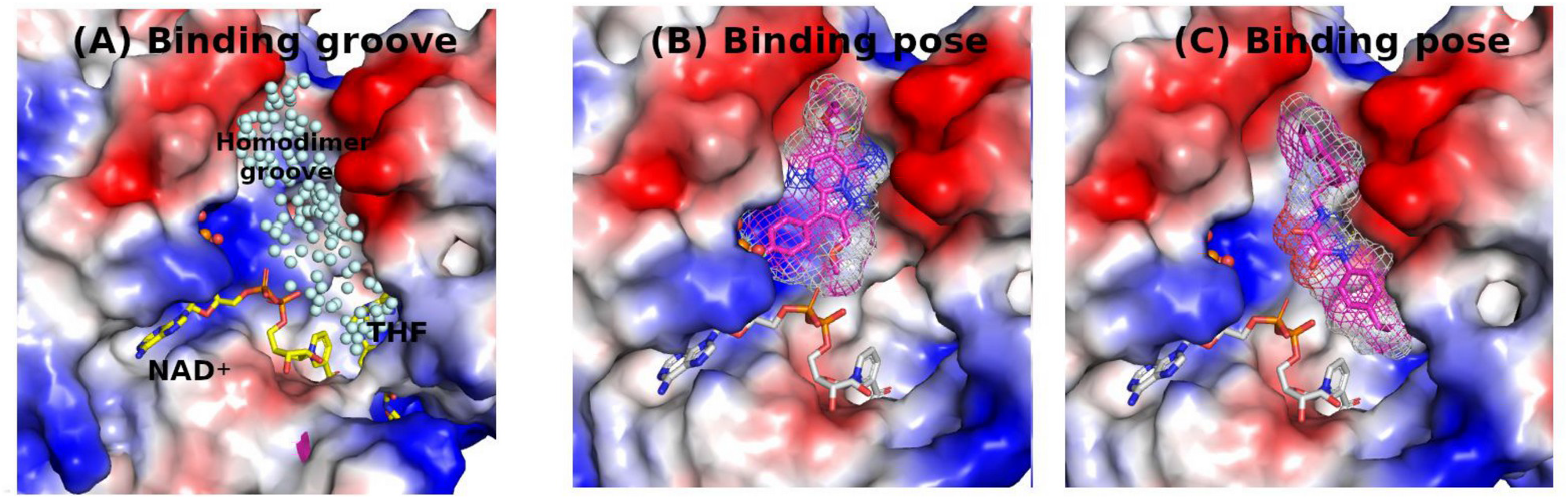
(A) Cofactor binding groove (NAD^+^), substrate groove (THF), and the homodimer groove integrated with a partial substrate groove which is filled with balls for illustration purposes. (B) and (C) are our preliminary screening poses for illustration purposes.

**Table S1:**
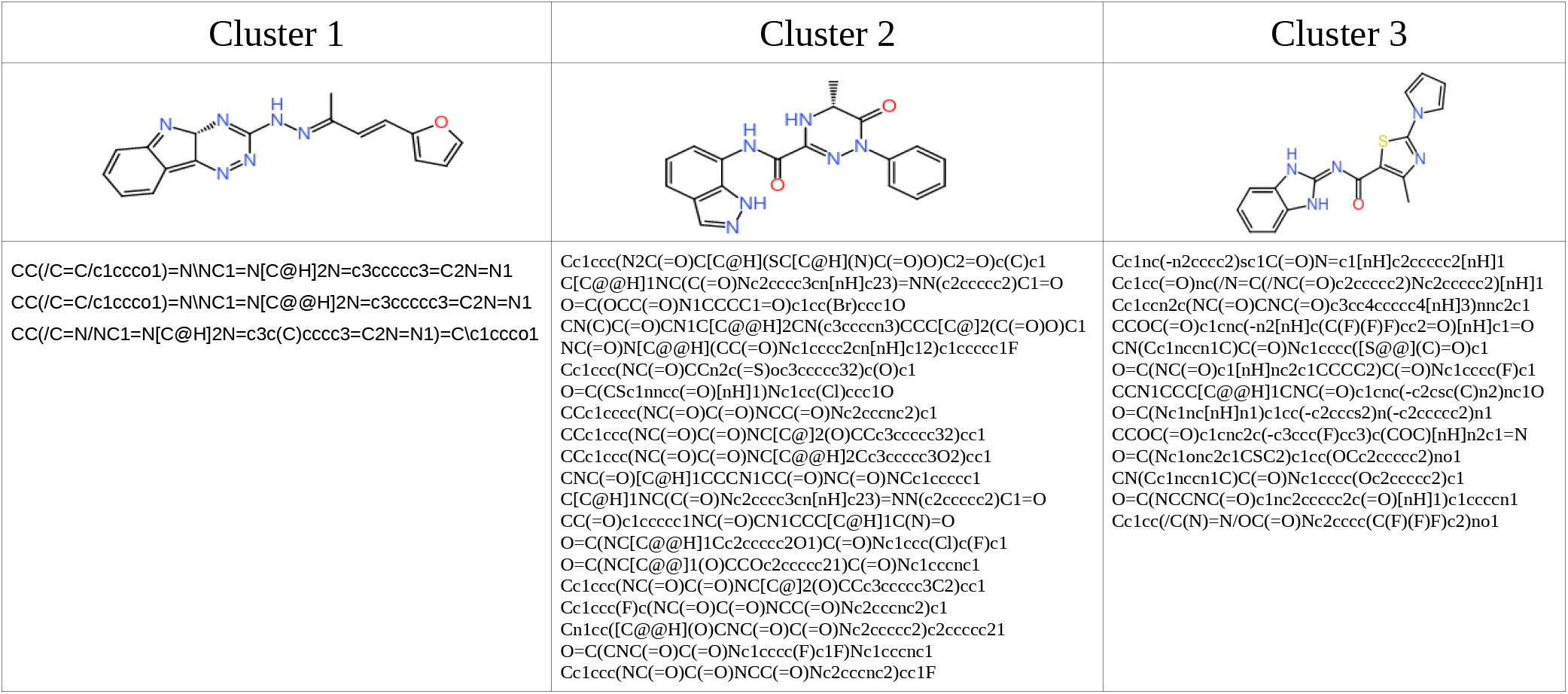
The compounds that bind the structurally less conserved homodimer groove (allosteric sites) from our preliminary screening.

### Empirical valence bond (EVB) simulations

The empirical valence bond (EVB) approach is a well developed and successfully used method.^2,3^ It allows you to practically explore the catalytic landscape by calculating reaction free energies in a practical and accurate way. It represents the reacting system, ranging from reactions of molecules to catalysis in enzymes, in a realistic but simple way. The system usually is represented by different resonance forms (diabatic states) with specifically curated force fields. Hence, it provides quantitative comparison of the effect of different environments (particularly in solutions and in enzymes) on the reaction potential surfaces. The resonance forms, namely the reactant, intermediate, and product states, are described by both the general molecular mechanics (MMs) force field within Enzymix for the region that is not involved in the reaction and a quantum empirical force field to represent the reaction region (EVB atoms). The complete protocol of EVB methods have been described well elsewhere,^2,4^ here we focus on the detailed parameters we used in our work.

**Figure S5:**
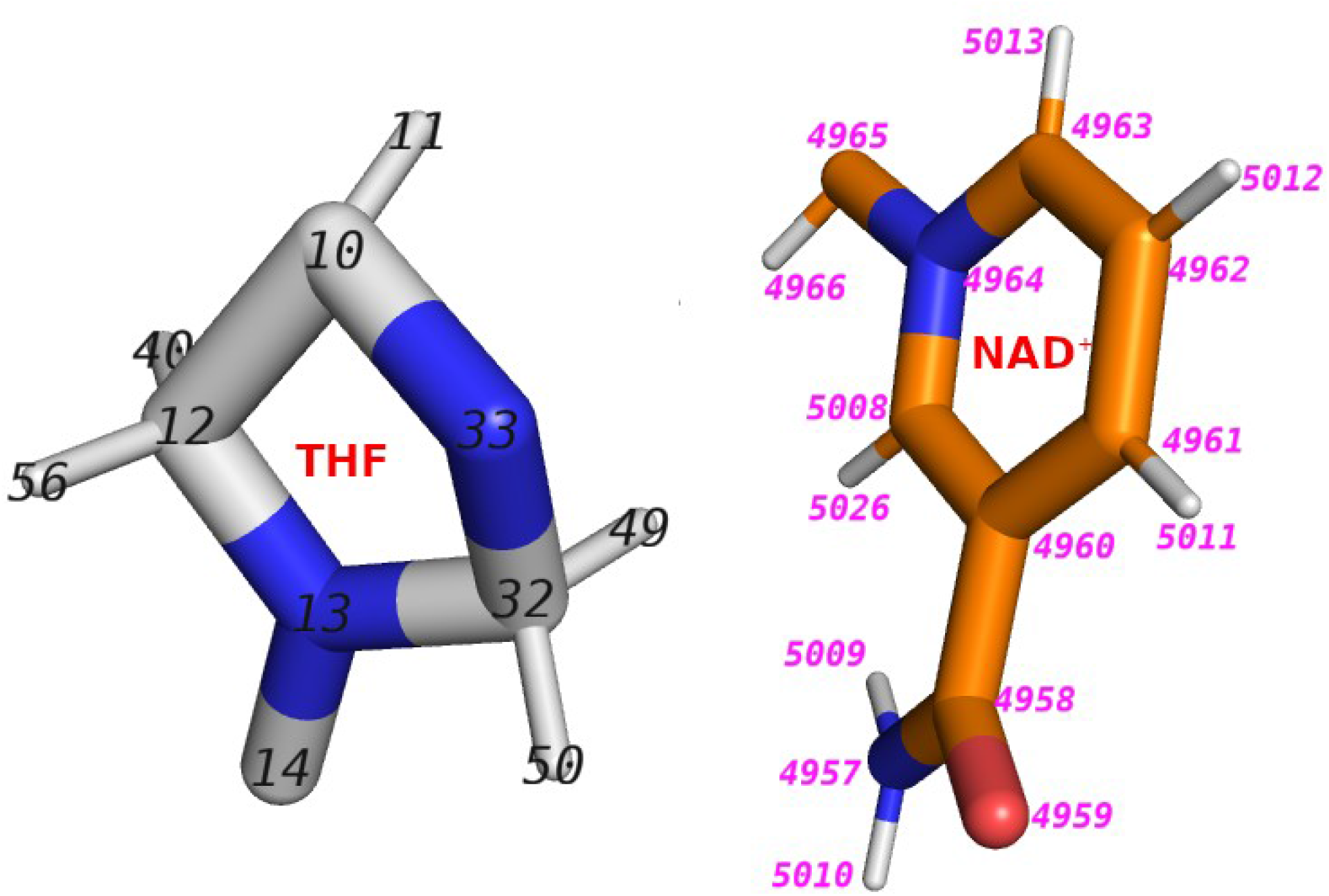
Depiction of the atom (in region I of our EVB simulations) with ID.

**Table S2:** Atom type and partial charges used in our EVB study.

| EVB atoms<br>(ID) | Reactant |  | Product |  |
| --- | --- | --- | --- | --- |
|  | Atom type | Charge | Atom type | Charge |
| 13 | N- | -0.324 | N- | -0.247 |
| 14 | C+ | 0.131 | C+ | 0.244 |
| 12 | C0 | 0.096 | C0 | 0.096 |
| 40 | H0 | -0.003 | H0 | 0.069 |
| 56 | H0 | 0.02 | H0 | 0.069 |
| 10 | C0 | 0.096 | C0 | 0.095 |
| 11 | H0 | 0.016 | H0 | 0.139 |
| 33 | N- | -0.339 | N- | -0.246 |
| 32 | C0 | 0.232 | C+ | 0.305 |
| 49 | H0 | -0.003 | H0 | 0.047 |
| 50 | H0 | -0.005 | H0 | 0.085 |
| 4957 | N- | -0.053 | N- | -0.088 |
| 4958 | C+ | 0.121 | C+ | 0.131 |
| 4959 | O0 | -0.271 | O0 | -0.328 |
| 4960 | C- | -0.114 | C- | -0.209 |
| 4961 | C+ | 0.206 | C0 | 0.117 |
| 4962 | C+ | -0.022 | C+ | -0.115 |
| 4963 | C+ | 0.204 | C+ | 0.120 |
| 4964 | N- | -0.155 | N- | -0.218 |
| 4965 | C0 | 0.071 | C0 | 0.076 |
| 5008 | C+ | 0.229 | C+ | 0.169 |
| 4966 | H0 | 0.273 | H0 | 0.146 |
| 5009 | H0 | 0.134 | H0 | 0.118 |
| 5010 | H0 | 0.129 | H0 | 0.112 |
| 5011 | H0 | 0.018 | H0 | 0.023 |
| 5012 | H0 | 0.04 | H0 | -0.012 |
| 5013 | H0 | 0.08 | H0 | 0.014 |
| 5026 | H0 | 0.071 | H0 | 0.015 |

**Table S3:** EVB bond parameters.

| Bonding interaction: $V_b = D_0[1 - e^{\mu(b-b_0)^2}] + k_b(b-b_0)^2$ | | | | | |
| --- | --- | --- | --- | --- | --- |
| Bond type | | $D_0(\text{kcal/mol})$ | $b_0(\text{\AA})$ | $\mu(\text{\AA}^{-1})$ | $k_b(\text{kcal/mol \AA}^{-2})$ |
| C0 | C0 | 96.0 | 1.54 | 0.8 | 400.0 |
| C0 | H0 | 100.4 | 1.10 | 0.8 | 400.0 |
| C0 | N- | 94.0 | 1.40 | 0.8 | 400.0 |
| C+ | C+ | 96.0 | 1.54 | 0.8 | 400.0 |
| C+ | C- | 96.0 | 1.54 | 0.8 | 400.0 |
| C+ | H0 | 100.4 | 1.10 | 2.0 | 400.0 |
| C+ | N- | 94.0 | 1.40 | 0.8 | 30.0 |
| C+ | O0 | 93.0 | 1.25 | 2.0 | 400.0 |
| N- | H0 | 98.3 | 1.10 | 2.0 | 30.0 |

**Table S4:** Lennard Jones Potential.

|  |  |  |
| --- | --- | --- |
| C+ | 632.00 | 24.00 |
| C- | 632.00 | 24.00 |
| C0 | 632.00 | 24.00 |
| H0 | 7.00 | 0.00 |
| N- | 774.00 | 24.00 |
| O0 | 774 | 24.00 |

**Figure S6:**
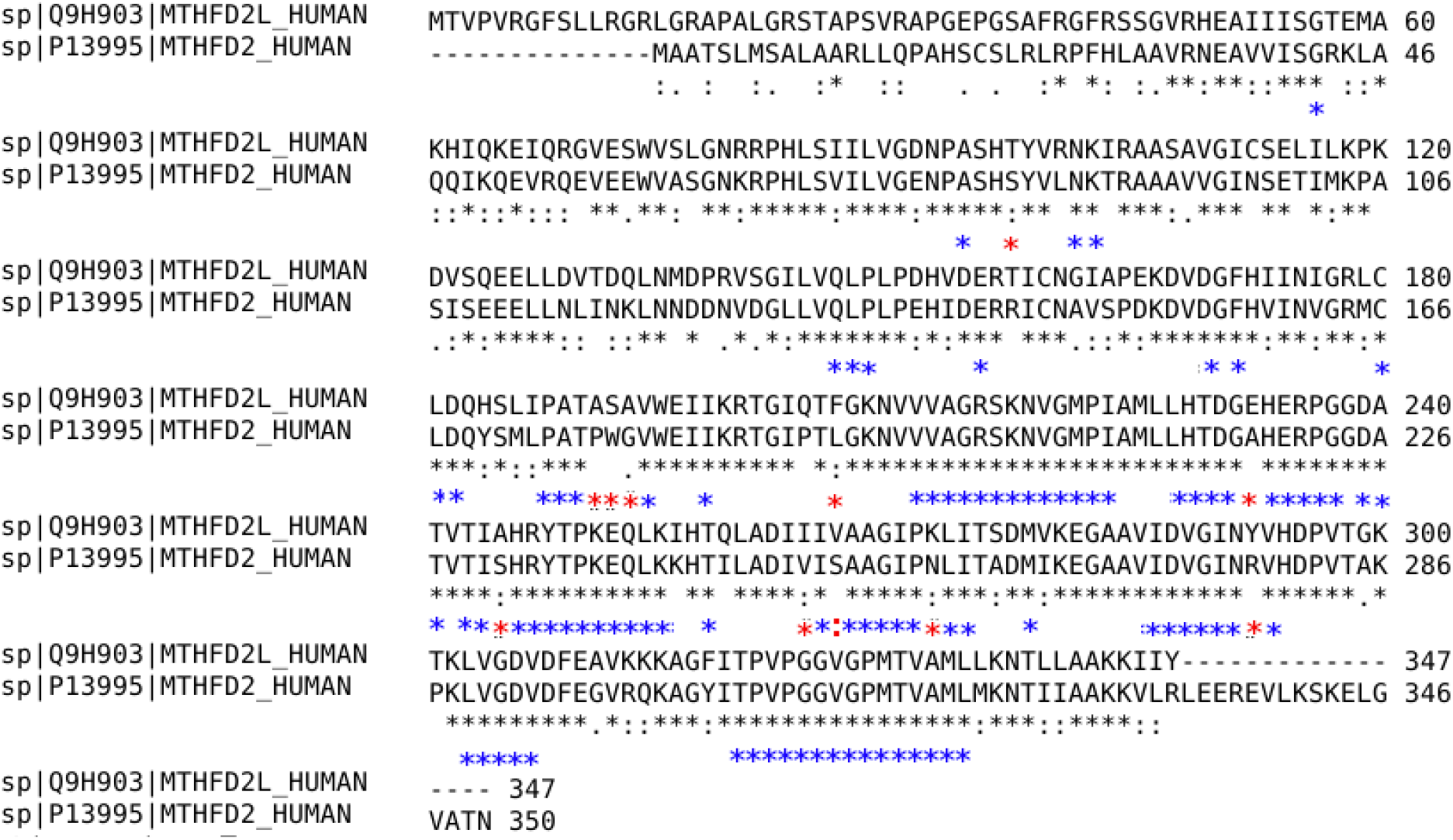
Sequence alignment of MTHFD2L and MTHFD2. The residues within 12Å from the NAD^+^ binding pocket were analyzed and colored blue (conserved between the two) and red (different between the two).

**Figure S7:**
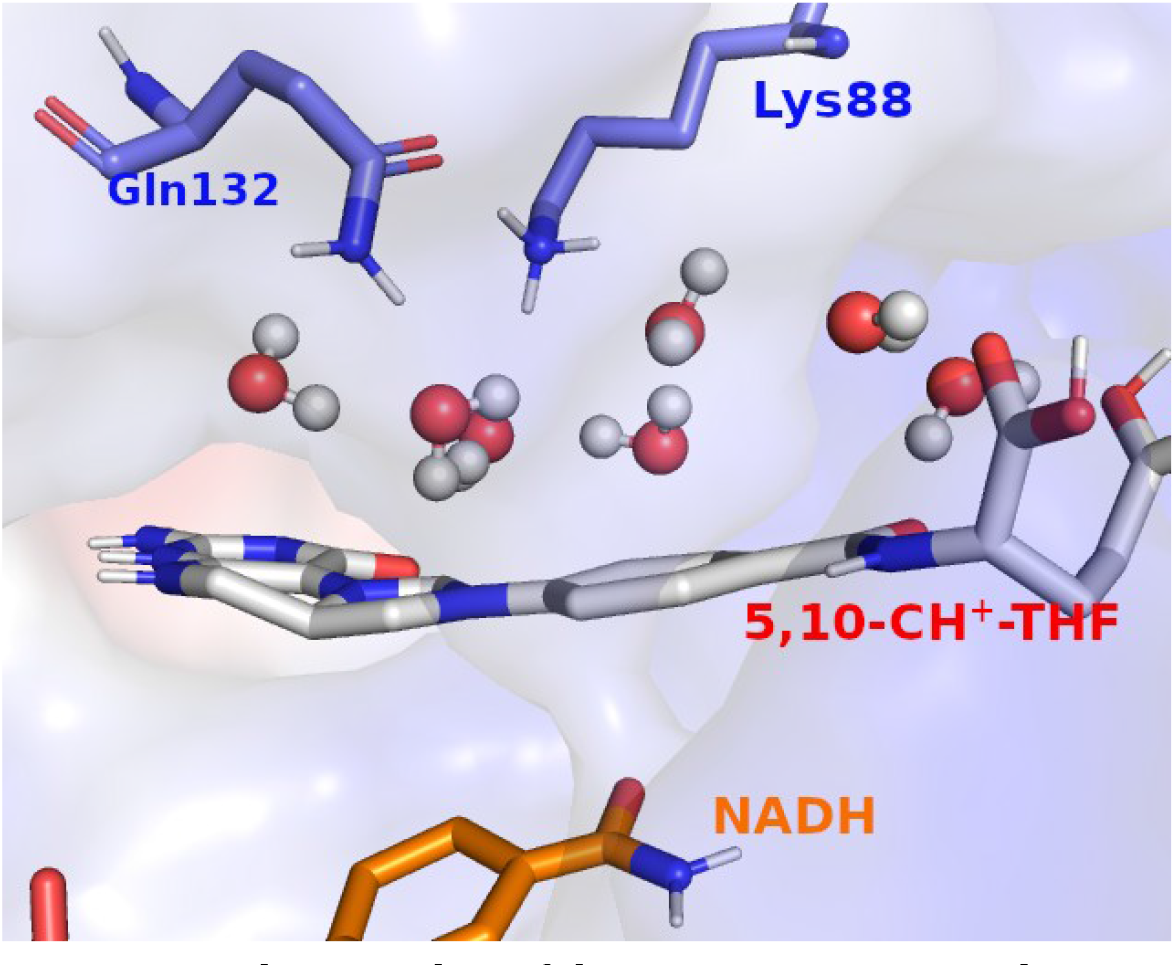
The snapshot of the protic proton-recycle-system.

**Figure S8:**
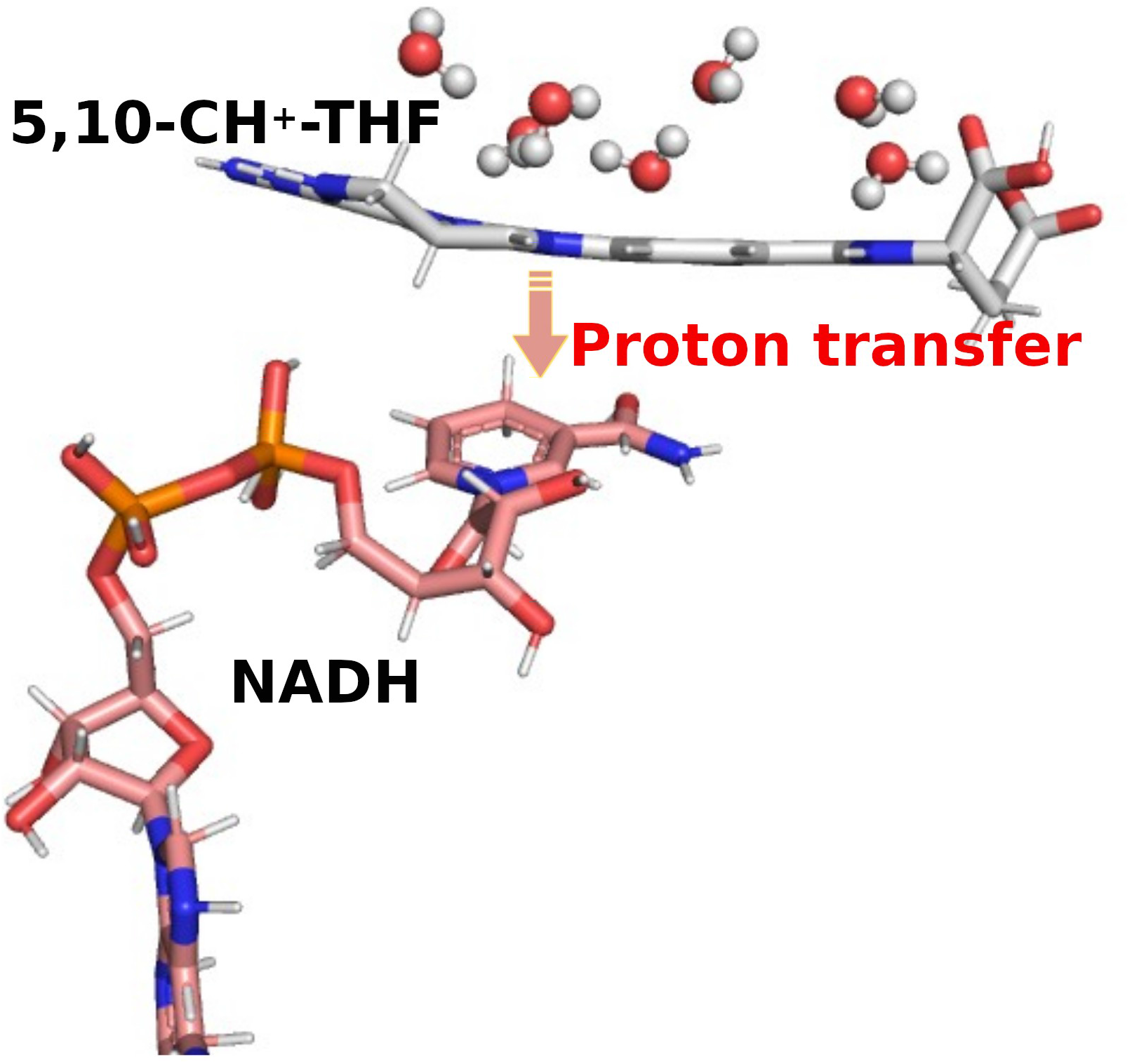
A snapshot to illustrate the catalytic process: the proton transfer from 5,10-CH_2_-THF to NAD^+^is accompanied by the charge redistribution with the dynamics of the water molecules featuring the re-ordering of water molecules to facilitate the subsequent cyclohydrolase reaction.

**Table S5:**
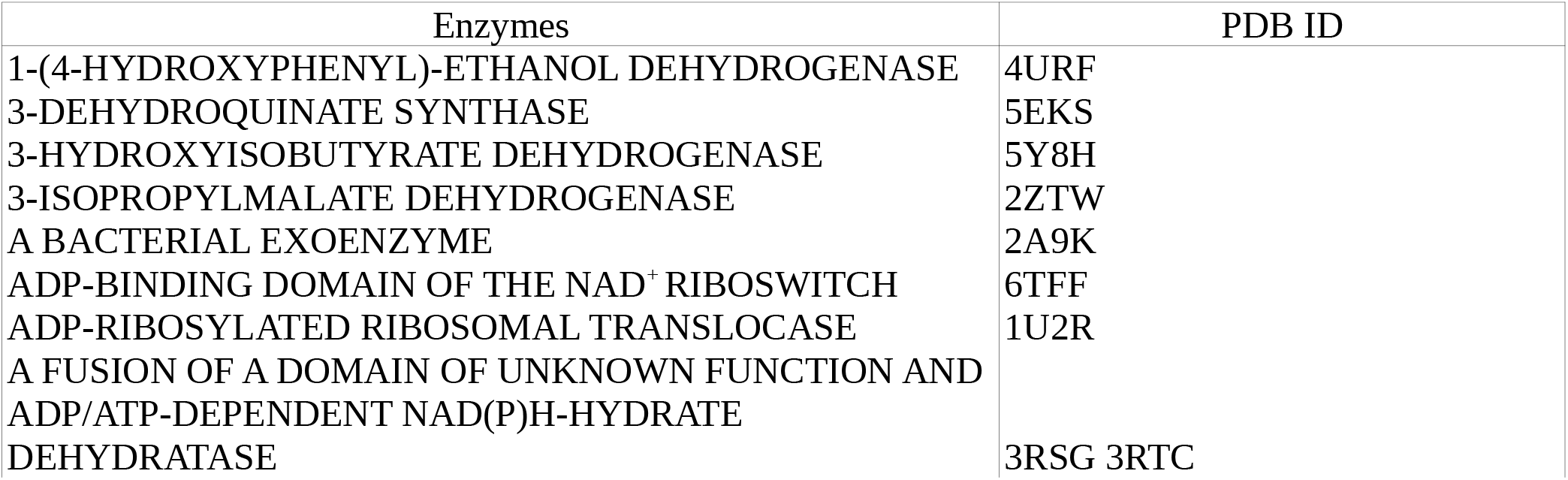
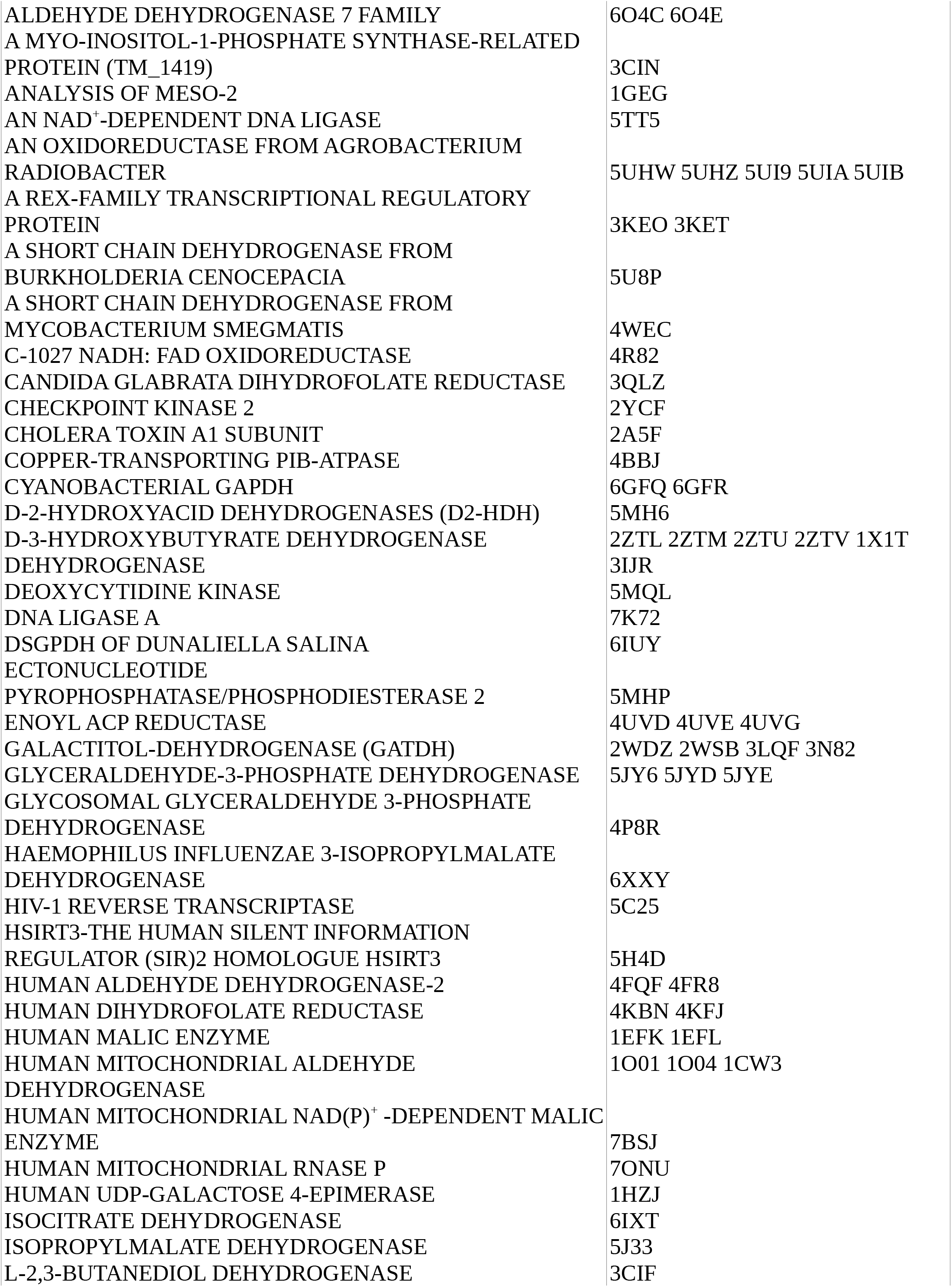
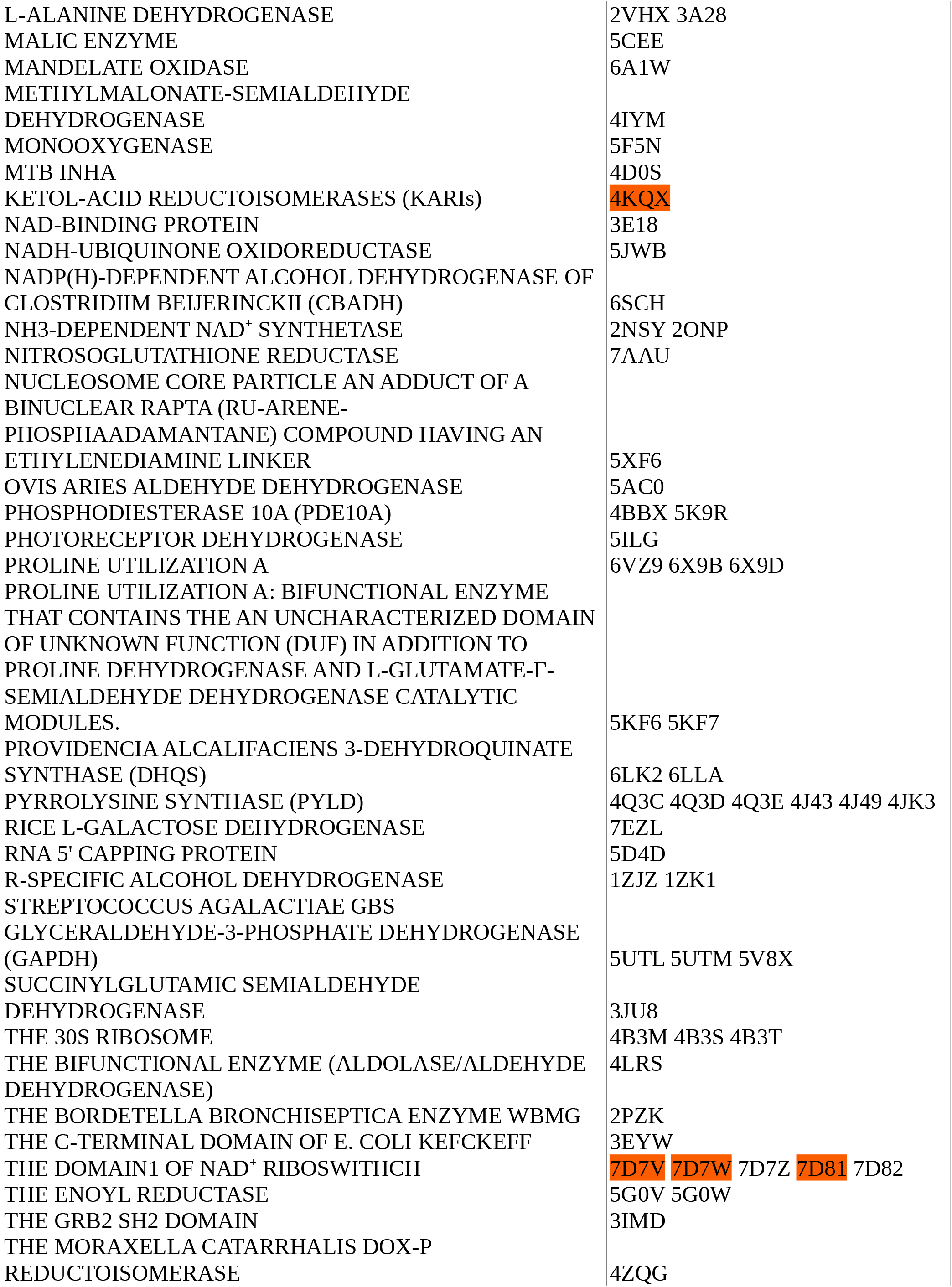
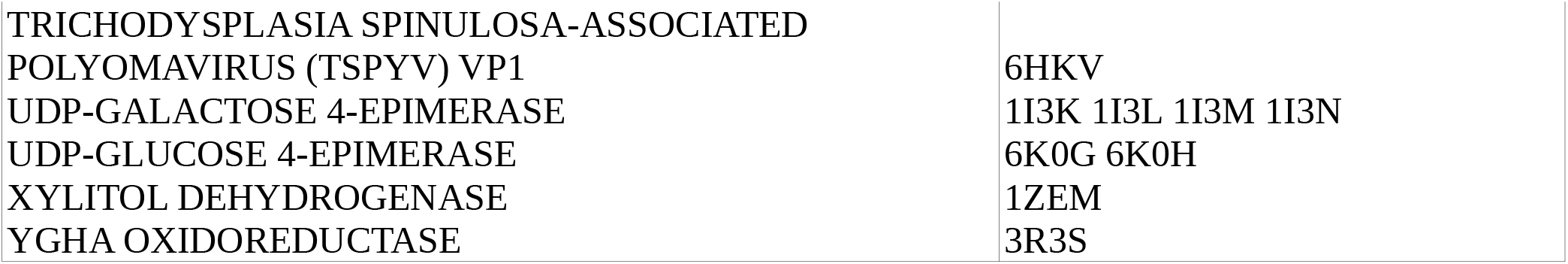
The proteins whose crystallized structures contain Mg^2+^ and/or NAD^+^. Note that the colored ones are the ones with two Mg^2+^ ions in close proximity of NAD^+^.

**Figure S9:**
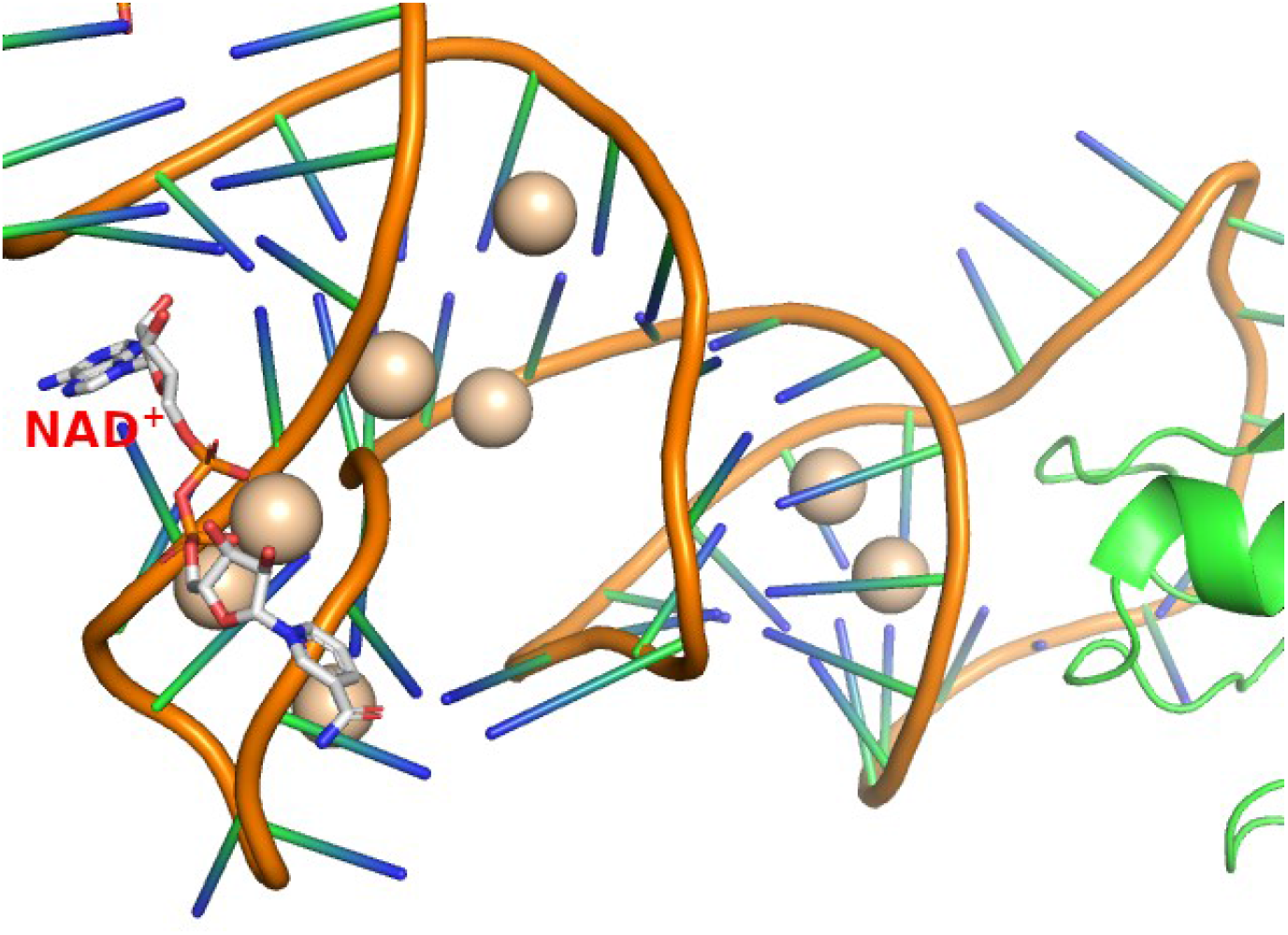
Snapshot to illustrate the NAD^+^ does not interact with protein from the structures of the riboswitch (PDB ID: 7D7V, 7D7W and 7D81). The NAD^+^ is shown as sticks while the Mg^2+^ are shown as spheres.

